# Temporal and spatial topography of cell proliferation in cancer

**DOI:** 10.1101/2021.05.16.443704

**Authors:** Giorgio Gaglia, Sheheryar Kabraji, Danae Argyropoulou, Yang Dai, Shu Wang, Johann Bergholz, Shannon Coy, Jia-Ren Lin, Rinath Jeselsohn, Otto Metzger, Eric P. Winer, Deborah A. Dillon, Jean J. Zhao, Peter K Sorger, Sandro Santagata

**Affiliations:** Laboratory of Systems Pharmacology, Department of Systems Biology, Harvard Medical School, Boston, MA, 02115, USA.; Ludwig Center at Harvard, Harvard Medical School, Boston, MA 02115, USA.; Department of Pathology, Brigham and Women’s Hospital, Harvard Medical School, Boston, MA 02115, USA.; Department of Medical Oncology, Dana Farber Cancer Institute, Boston, MA, 02215, USA.; Department of Cancer Biology, Dana Farber Cancer Institute, Boston, MA, 02215, USA.; Harvard Graduate Program in Biophysics, Harvard University, Cambridge MA 02138, USA.; Department of Medicine, Brigham and Women’s Hospital and Harvard Medical School, Boston, MA 02115, USA.; Department of Oncologic Pathology, Dana Farber Cancer Institute, Boston, MA, 02215, USA.

**Keywords:** proliferation, cell cycle, multiplexed imaging, cancer systems biology, temporal inference, spatial architecture, CyCIF, glioblastoma, mesothelioma, breast cancer

## Abstract

Proliferation is a fundamental trait of cancer cells but is poorly characterized in tumors by classical histologic methods. We use multiplexed tissue imaging to quantify the abundance of multiple cell cycle regulating proteins at single-cell level and develop robust multivariate proliferation metrics. Across cancers, the proliferative architecture is organized at two distinct spatial scales: large domains, and local niches enriched for specific immune lineages. A subset of tumor cells express cell cycle regulators in canonical patterns consistent with unrestrained proliferation, a phenomenon we refer to as “cell cycle coherence”. By contrast, the cell cycles of other tumor cell populations are skewed toward a specific phase or characterized by non-canonical (incoherent) marker combinations. Coherence varies across space, with changes in oncogene activity, and with therapeutic intervention, and is associated with aggressive behavior. Multivariate measures capture clinically significant features of cancer proliferation, a fundamental step in enabling more precise use of anti-cancer therapies.

## INTRODUCTION

Although uncontrolled cell proliferation is a defining feature of cancer (Hanahan and Weinberg, 2011) much of our understanding of the cell cycle comes from the *in vitro* study of cell monocultures grown in an abundance of nutrients, supporting doubling times of 24 to 48 hours (Duval et al., 2017). By contrast, in patients the median doubling time of tumors is substantially longer; even aggressive tumors such as colorectal metastases to the lung have a median doubling time of over 90 days (Collins et al., 1956). While it has been long appreciated that most solid tumors do not uniformly grow according to exponential kinetics (Norton, 1988; Norton et al., 1976), the molecular and cellular determinants of proliferation in the microenvironment of a tumor are incompletely understood (Norton, 2014).

While mutations in oncogenes and tumor suppressors are a prerequisite for cancer growth, cell-intrinsic growth signals arising from these genetic changes are only one component of a rich network that influences whether or not cancer cells divide in vivo (Black and McGranahan, 2021). The levels of nutrients, oxygen, and metabolites, as well as the physical properties of the tumor (Nia et al., 2020) vary dramatically between different tumors and spatially within single lesions, each potentially imposing restrictions on cell proliferation (Frieboes et al., 2006; Randall et al., 2020). Thus, in both primary and metastatic tumors, cancer cells exist in proliferative, non-proliferative, and arrested states (Aguirre-Ghiso, 2007). In addition, many cells in tumor masses are not neoplastic, but rather are immune and stromal cells (Quail and Joyce, 2013) that are non-uniformly distributed across the tumor (Lin et al., 2021). The extensive crosstalk between these distinct cellular components of a tumor ecosystem creates local conditions that may either promote or inhibit cell proliferation (Bejarano et al., 2021).

Tumors that are removed during clinical care contain valuable information about the tumor cell proliferation and how it is influenced by resource limitation and the physical constraints imposed by surrounding tissues. In the past, autoradiography measurements of ^3^H-thymidine incorporation into DNA from resected tumors provided information about cell cycle kinetics and demonstrated that only a fraction of cells were actively proliferating (Baserga, 1965; Johnson et al., 1960; Mauer and Fisher, 1962). Today, in both research and clinical diagnosis, tumor cell proliferation is primarily assessed using two features detectable by tissue imaging: the frequency of mitotic figures as judged visually in hematoxylin and eosin (H&E) stained tissue sections or the fraction of Ki-67-positive cells as measured using immunohistochemistry (IHC) (Inwald et al., 2013). Both measures have substantial limitations. Mitotic figures reflect one brief phase of the cell cycle, and do not always represent active proliferation as they can also accumulate during mitotic arrest; their detection is also highly dependent on staining and fixation quality (Lehr et al., 2013). Ki-67 is not an essential cell cycle regulator but rather a protein that organizes chromatin during mitosis and whose levels correlate with proliferation (Cuylen-Haering et al., 2020; Sobecki et al., 2016). Studies in cultured cells show that Ki-67 levels change in a graded manner throughout the cell cycle, rising gradually during S phase, peaking during mitosis, and falling during anaphase and G1 (Bruno and Darzynkiewicz, 1992; Miller et al., 2018). Nonetheless, in clinical practice the proliferative index of tumors is scored as the percentage of Ki-67 positive cells, requiring each cell to be assigned a dichotomous score as Ki-67 positive (proliferating) or Ki-67 negative (non-proliferating). The imprecision of this approach underestimates the proportion of cells that are actually proliferating (Gerdes et al., 1983; Miller et al., 2018). Multiple studies have demonstrated the potential of proliferative index to serve as both a prognostic and predictive biomarker, but the analytic and pre-analytic variability of Ki-67 staining has made it difficult to realize this promise (Denkert et al., 2015; Nielsen et al., 2020). A more accurate and comprehensive means to assay proliferation that accounts for the complexities of cell cycle dynamics is therefore essential for applications as diverse as basic and translational research, clinical trials, patient management, and tissue and tumor atlas construction (HuBMAP Consortium, 2019; Rajewsky et al., 2020; Rozenblatt-Rosen et al., 2020).

The multiplexed measurements necessary to deeply characterize cell proliferation in clinical specimens have only recently become possible. Over the past several years, a range of multiplexed tissue imaging methods have been introduced to deeply phenotype fixed tissues (Angelo et al., 2014; Giesen et al., 2014; Goltsev et al., 2018; Lin et al., 2018; Saka et al., 2019; Tsujikawa et al., 2017). These methods measure the levels of 10-60 antigens at single cell resolution and permit the identification and quantification of cell types and cell states as well as the description of cell-cell interactions and higher-order relationships in space (Bodenmiller, 2016). Collaborative projects such as the NCI Human Tumor Atlas Network (HTAN) are using these technologies to create spatial maps of human cancer in which architectural and cell state features are related to clinical outcomes (Rozenblatt-Rosen et al., 2020). Multiplexed protein imaging is well-suited to studying processes such as cell cycle progression, which are regulated by oscillatory proteolysis of cell cycle phase-specific proteins, and can therefore be monitored using protein level measurement alone (Gookin et al., 2017; Mahdessian et al., 2021). For example, the inverse oscillations of DNA licensing factors CDT1 (Nishitani et al., 2000) and Geminin (McGarry and Kirschner, 1998) through G1/S/G2 have been used to delineate cell cycle phase transitions in vitro and in vivo (Sakaue-Sawano et al., 2008). Imaging at sub-cellular resolution makes it possible to quantify the translocation of proteins between cellular compartments, as well as the breakdown in nuclear structure at mitosis. The translocation of cyclin B1 from the cytoplasm to nucleus, for example, is a reliable way to monitor the G2 to M transition in mammalian cells (Jin et al., 1998). One complication in this approach is that cancers frequently carry mutations or copy number changes in cell cycle regulators, such as *RB1* (pRB) and *CKDN2A* (p16) (Priestley et al., 2019; Robinson et al., 2017). This causes disruption of normal cell cycle progression and impacts protein expression patterns. For example, loss of pRB results in hyperexpression of p16 as feedback mechanisms unsuccessfully attempt to block aberrant cell cycle progression (Romagosa et al., 2011; Shapiro et al., 1995). With information on many different proteins provided by multiplexed measurements, single cell patterns of correlation and decorrelation among cell cycle regulators can be probed for insight into the fidelity of cell division. Multiplexed single cell measurements also reveal connections between the levels of expression or activities of oncogenic proteins (e.g., the HER2 receptor in breast cancer) and cell cycle dynamics (Wolf-Yadlin et al., 2006).

In this study we use multiplexed measurements of cell cycle regulators from fixed tissues in two ways. First, we develop a multivariate proliferation index (MPI) that incorporates information from multiple markers to categorize tumor cells as proliferative, non-proliferative or arrested. This corrects for biases arising from the use of Ki-67 alone as a measure of proliferation. Second, we create a framework for studying cell cycle dynamics from fixed cell images based on time inference, a computational method to model dynamic processes in the absence of temporal data. Several time inference methods have been developed for inferring dynamics from fixed time images acquired from cells grown in culture (Gut et al., 2015; Kafri et al., 2013) and from single-cell RNA sequencing data (Cannoodt et al., 2016; Liang et al., 2020; Setty et al., 2019; Trapnell et al., 2014). To develop an analogous approach for multiplexed images acquired from fixed tumor tissues we made the following assumptions: 1) measured markers provide coverage of multiple cell cycle transitions, 2) sampling is sufficiently uniform and dense for ergodic theory to be applicable, and 3) there exists a dynamical system (not necessarily known) governing changes in expression of measured cell cycle proteins. Similar assumptions are needed to apply any time inference framework to static data. When these assumptions are met, the time inference method approximates time series data and can be used to order cells in ‘pseudotime.’

Using these two approaches, we address fundamental questions about cell proliferation in cancer tissues. We identify general properties about the organization of proliferating cells across major tumor types and demonstrate short-range tightly organized niches and long-range zones having graded features. Locally, proliferative and non-proliferative niches are enriched for distinct types of immune cells. Using an interpretable visualization of the multidimensional space of cell cycle markers we identify three distinct types of cell cycle dynamics: coherent, skewed, and non-canonical. Coherent dynamics in tissues reflect the pattern seen in freely cycling cells grown in culture. Studies in HER2 breast cancer tissues reveal an unexpected relationship between oncogene expression levels and cell cycle progression: optimal coherence is observed at intermediate HER2 levels, not with the highest receptor expression, consistent with the existence of optimum oncogene expression. In patient samples acquired before, on, and after treatment, adaptive changes in cell cycle dynamics are observed, showing that imaging cell cycle regulatory proteins can be applied to understand the effects of anti-cancer drugs and to potentially guide their use. Finally, we show that cell cycle coherence is associated with tumor recurrence in two highly aggressive malignancies, glioblastoma and mesothelioma. Thus, our results show how the correlation structure of multi-parameter single cell protein expression data can be used to study temporal processes that can only be assayed in humans at a single point in time. When combined with the spatial and morphological data available in images, this provides a new means to investigate the complexities of human disease.

## RESULTS

### A Multivariate Proliferation Index (MPI) derived from multiplexed immunofluorescence images

We used 20 to 30-plex cyclic immunofluorescence (CyCIF) to image sections from formalin fixed paraffin embedded (FFPE) human epithelial cancers (i.e., breast, lung, colon, ovarian carcinomas), mesothelioma, and gliomas, in each case recording the intensities of each immunofluorescence signal on a per-cell basis (∼21.3 million cells in total, from >500 individual specimens; **Table S1**). We used lineage-specific markers (e.g., e-cadherin, pan-cytokeratin, SOX2, CD45, vimentin), to distinguish tumor cells from immune and stromal cells and characterized the proliferative states of the tumor cells (**Figures 1-2**). We then investigated cell cycle properties in multi-dimensional marker space, before and after perturbation, by interpreting marker combinations in light of known protein expression patterns across the cell cycle (**Figures 3-7**).

**Figure 1.**
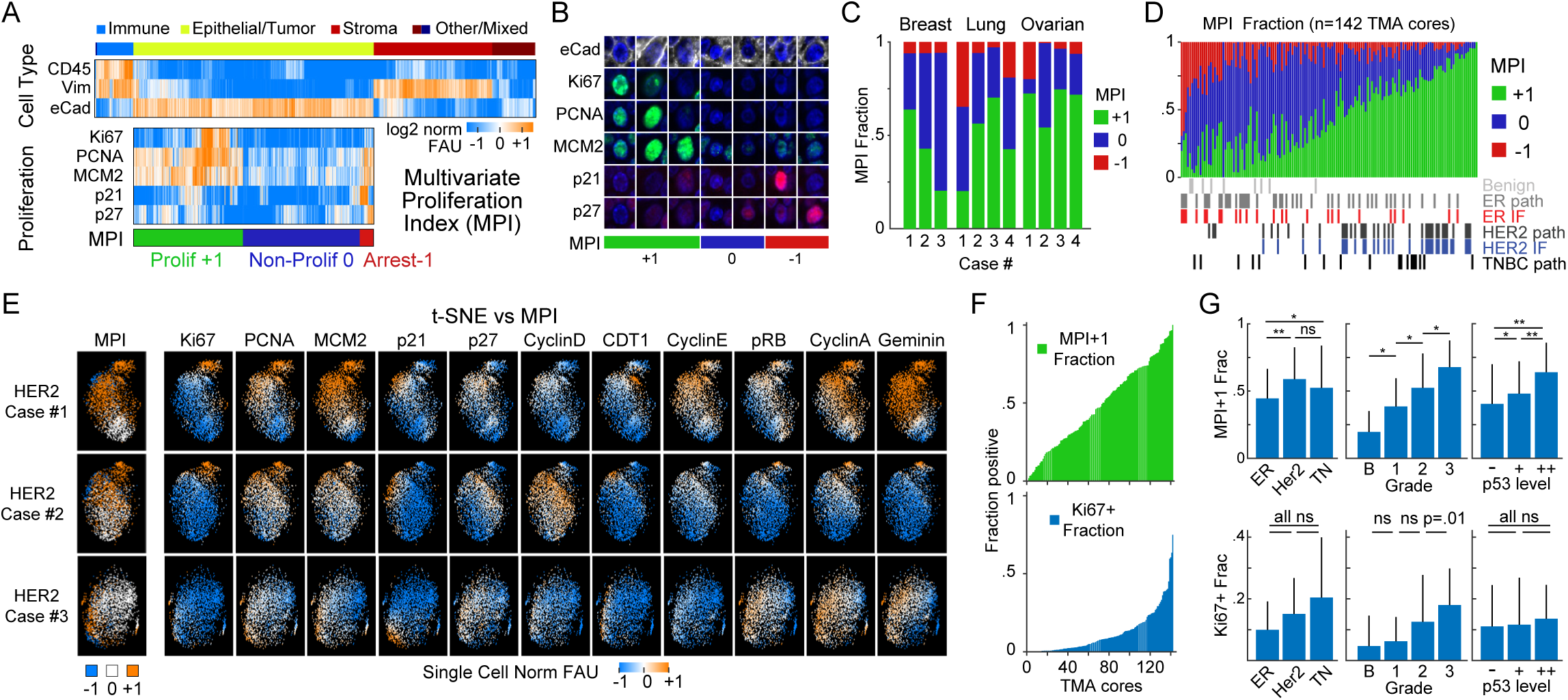
Multivariate Proliferation Index (MPI) is linked to clinical parameters. (A) Top, clustered heat map of normalized log2 fluorescence signal intensities of cell lineage markers from CyCIF images (one cell per column, breast tumor tissues n = 2.5 million cells). Bottom, clustered heat map of signal intensities of five markers in epithelial/tumor population (n = 1.4 million cells), sorted by MPI categories: +1 (proliferative, green), 0 (non-proliferative, blue), or -1 (arrested, red). (B) Representative immunofluorescence images of individual tumor cells from breast cancer showing MPI marker expression and corresponding MPI category. (C) Stacked bar graph of epithelial/tumor cells of MPI categories from samples of tissues from three types of carcinomas (n = 3 breast, 4 lung and 4 ovarian cases) (D) Stacked bar graph of epithelial/tumor cells of MPI categories from 142 samples of tissues from 75 patients (Pantomics BRC15010). The receptor status for each tumor is indicated according to the vendor (‘path’) and from direct CyCIF measurements. (E) t-SNE plots for the three breast carcinoma tissues from panel C. with proliferation and cell cycle markers mapped to color (MPI categories were not used as t-SNE variables, n = 2,500 cells). (F) Comparison between MPI +1 fraction and Ki-67+ fraction per core in breast tissue microarray (n = 142 cores from panel D, ordered independently by metric). (G) Comparison of MPI+1 and Ki-67 positive fraction in epithelial/tumor cells across different classifiers of breast cancer (n = 142 cases from panel D, KS p-values *p<0.035, ** p<0.006).

As a robust means to aggregate information on cell proliferation we generated a Multivariate Proliferation Index (MPI). This categorical index is based on staining intensities for three proliferation markers (Ki-67, PCNA, MCM2) (Bravo et al., 1987; Chong et al., 1995; Madine et al., 1995; Takasaki et al., 1981) and two cell cycle arrest markers (p21, p27) (Cayrol et al., 1998; Sherr and Roberts, 1999) (**Figures 1A, 1B**, and **S1A-S1D**). An MPI value was assigned to each tumor cell based on the following rule: cells were scored as *proliferative* (MPI +1) if they expressed a positive balance of proliferation markers; *non-proliferative* (MPI 0) if they lacked expression of proliferation markers; and *arrested* (MPI -1) if they expressed high levels of one or both of the arrest markers, even if proliferation markers were also expressed. The frequency of proliferative cells (MPI +1) varied between tumor samples (**Figures 1C** and **1D**), but MPI calculations from adjacent sections of the same samples were reproducible, demonstrating technical robustness (linear fit coefficient = 1.004, R^2^ = 0.89; **Figure S1E**). No single marker appeared sufficient for identifying all proliferative cells (**Figure 1 A,B**; **Note S1**). For example, although Ki-67 is the most widely used measure of proliferation in diagnostic and research settings (Allegra et al., 2003; Viale et al., 2008), we found that 39-72% of MPI +1 cells were Ki-67 negative but positive for PCNA or MCM2, depending on the tumor type; this is consistent with data from cultured cells showing that high Ki-67 expression occurs in the G2 phase of the cell cycle (Bruno and Darzynkiewicz, 1992; Miller et al., 2018) (**Figures S1B** and **S1C**). Results from MPI classification were consistent with those obtained by dimensionality reduction followed by clustering of single cell data (e.g., with t-SNE in **Figure 1E;** and with k-means in **Figure S1D**), but computing MPI is advantageous because supervised labelling and parameter tuning are not required to identify the proliferative subpopulations (**Note S2**). MPI is likely to be a more reliable measure of proliferation than Ki-67 staining alone because MPI encompasses multiple cell cycle states (e.g., G1/S and G2/M) and involves redundant measurements, making it less sensitive to staining artifacts.

To determine whether the variation in MPI across patient samples is associated with clinically relevant features of tumor behavior, we focused on breast cancer in which tumor subtype (DeSantis et al., 2019), histological grade (Rakha et al., 2010), and p53 status (Allred et al., 1993) are known predictors of outcome. In samples from 75 patients, the fraction of MPI +1 cells varied from 0 to 1 (**Figure 1F**) and was highest in aggressive molecular subtypes such as HER2-amplified and triple-negative breast cancer (KS test, p < 0.035), **Figure 1G**). The fraction of cells scored as MPI +1 also increased significantly with tumor grade and p53 status (KS test, p-value < 0.035, **Figures 1D** and **S1F**), but this was not true of the Ki-67 positive fraction (**Figure 1G**).

### Spatial analysis of MPI reveals two length scales of proliferative architecture of human cancer

To determine whether proliferating and non-proliferating tumor cells are organized into distinct spatial domains, we quantified the spatial correlation within MPI categories (“self-correlation”) and between MPI categories (“cross-correlation”) across 513 tumor specimens including carcinomas, mesothelioma and gliomas. Visual inspection of multiplexed image data and corresponding single cell maps of MPI values revealed a variety of spatial patterns (**Figures 2A** and **S2A-S2C**). Cross-correlations were found to be weak and variable, but proliferative (MPI +1) and non-proliferative (MPI 0) states were strongly and significantly positively self-correlated across tumor types (**Figures 2B** and **S2D-S2F**). Spatial self-correlation decreased with distance and was well fit by a two-exponential decay model (**Figures 2C** and **S2G**). From the fitting, we estimated two characteristic length scales corresponding to ∼10-30 µm and ∼100-300 µm (**Figures 2D**). Thus, proliferating (MPI +1) cells are clustered together with other proliferating cells and away from non-proliferating and arrested cells. Further, the proliferative architecture is organized in two physical scales; small niches within larger structured neighborhoods.

**Figure 2.**
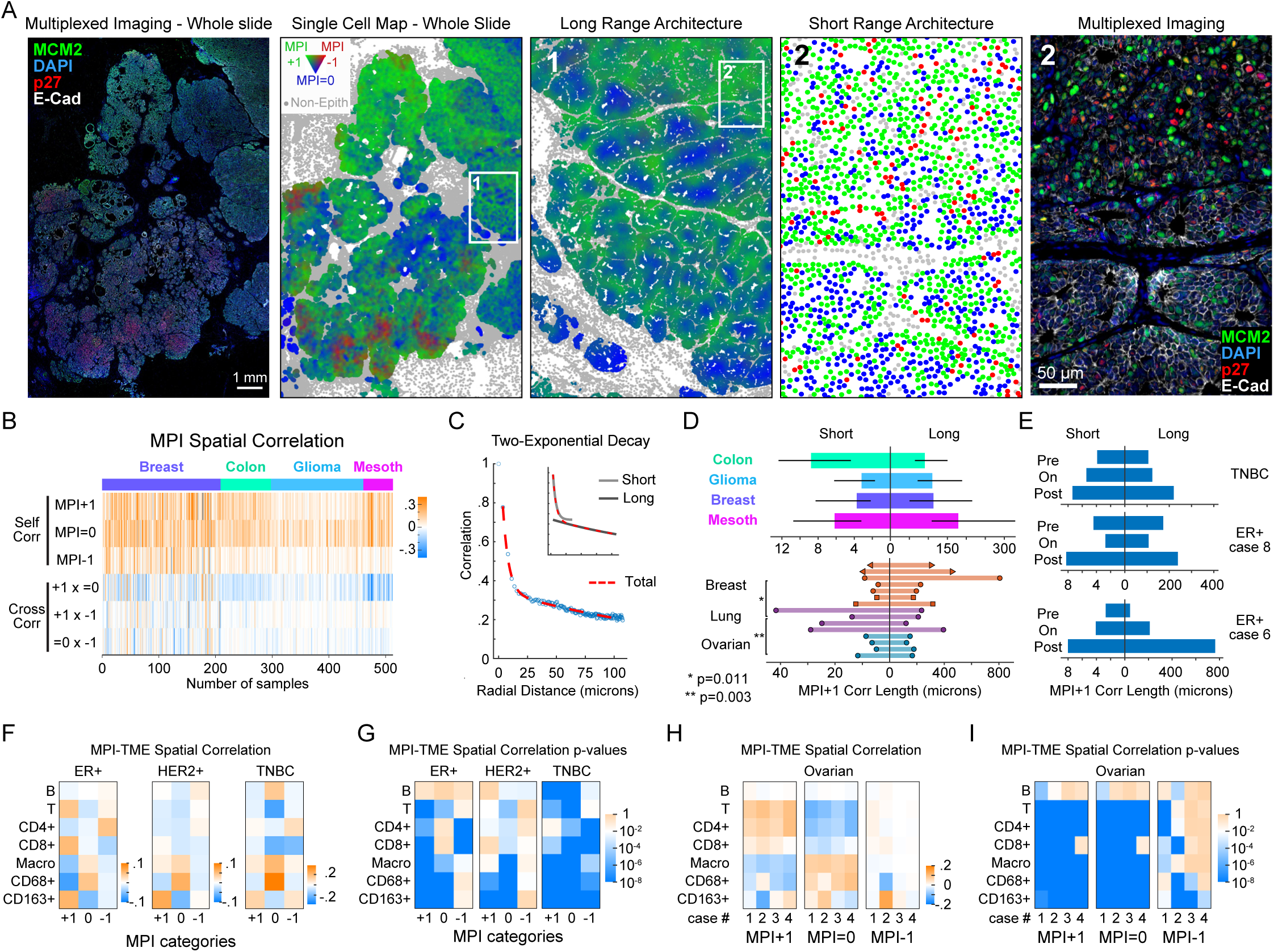
MPI reveals two proliferative domains of cancer proliferative architecture. (A) Spatial maps of MPI categories (whole slide and inset 1 were smoothed over 40 neighboring cells, for visualization purposes only) and corresponding composite CyCIF images from a HER2-positive breast tumor (white = E-Cadherin, green = MCM2, red = p27, blue = DNA). (B) Heat map of spatial correlations within and across MPI categories (“self corr” and “cross corr” respectively, k = 5^th^ neighbor, n = 513 samples). (C) Spatial correlation plot and two-exponential fit. Inset depicts the two exponential curves that composed the fit (“short “and “long” scales). (D) Spatial correlation decay lengths for four tissue microarrays (median +/- 25th percentile, 53 breast, 73 colon, 122 glioma, 32 mesothelioma samples) and 15 whole slide cancer tissues (7 breast, 4 lung and 4 ovarian). (E) Spatial correlation lengths through treatment (see **Table S5** for treatment details). (F-I) Spatial correlation between epithelial tumor cells and immune cells and corresponding p-values for (F-G) breast sample cohort (ER+ n = 46, HER2+ n = 37, TNBC n = 18 samples, Pantomics BRC15010) and (H-I) individual ovarian whole tumor slides (n = 4). Pearson correlation p-values are displayed in log10 color scale.

When we calculated characteristic length scales for MPI data in tissues from three breast cancer patients biopsied before, on and after treatment, both short and long correlation lengths increased following therapy (**Figure 2E**). At short length scales in breast tissue from 75 patients (from **Figures 1D, 1F and 1G** above), MPI 0 tumor cells clustered away from T lymphocytes and were more frequently found in proximity to CD68-expressing macrophages, whereas MPI +1 tumor cells were associated with CD163-expressing macrophages (correlation p-value <10^-8^) (**Figures 2F** and **2G**). In ER+ breast cancer, MPI +1 cells were significantly associated with cytotoxic T cells (correlation p-value <10^-8^) (**Figures 2F** and **2G**); this strong association was also observed in ovarian cancer (**Figures 2H** and **2I**) but was absent in HER2+ and triple negative breast cancers (**Figures 2F** and **2G**). We speculate that the short length scales observed in MPI data correspond to small clusters of sister cells arising from common parents that are also influenced by interaction with immune cells. Large proliferative neighborhoods may be organized by environmental conditions e.g., hypoxia (Tannock, 1968; Zaidi et al., 2019), nutrient availability, and tissue structure (Muthuswamy, 2021).

### Cell cycle coherence metrics derived from multiplexed images of human cancer

In addition to allowing us to visualize the spatial distribution of proliferation in cancer, MPI identified actively proliferating cells for further characterization. We therefore stained tissues using antibodies against 10 well-established cell cycle regulators including cyclins A1/A2, B1, D1 and E1, CDK inhibitors p21 and p27 and DNA replication regulators CDT1, Geminin, and phospho-RB (see **Table S2** for details on markers used in each experiment). Single cell intensity data revealed a wide range of marker combinations (**Figures 3A** and **S3A**), but most markers did not have clearly separated high and low states (**Figures 3B** and **S3B-S3C**). Neither multidimensional gating nor dimensionality reduction methods such as t-SNE provided insight into the likely order of cell cycle events (**Figures 1E** and **S3D**; **Notes S2-S4**). We therefore sought an alternative approach informed by knowledge of cell cycle dynamics.

**Figure 3.**
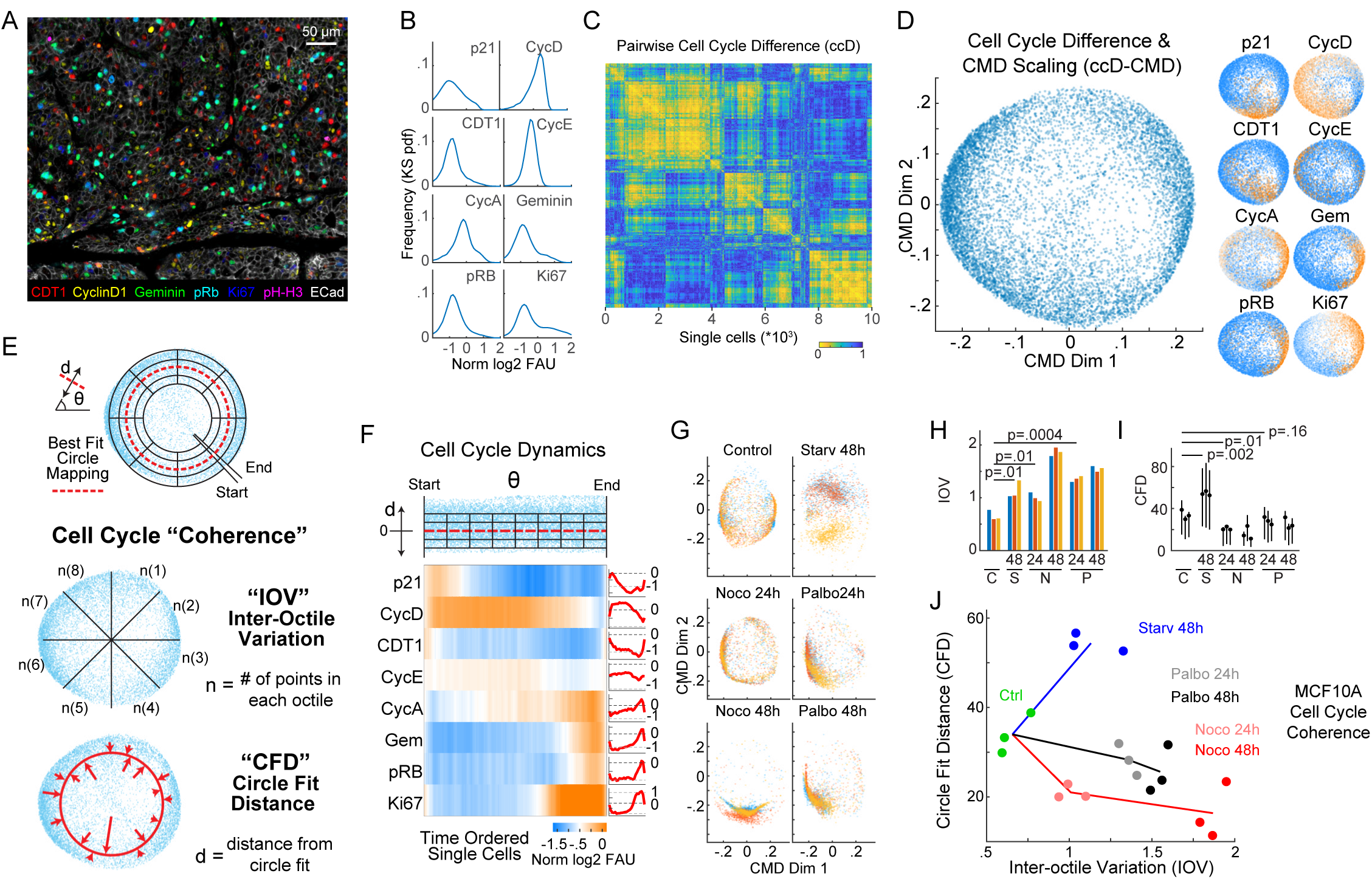
A framework for inferring cell cycle dynamics from multiplexed imaging data. (A) Composite fluorescence image of a subset of cell cycle markers and e-cadherin (breast tumor, scalebar, 50 µm). (B-E) Computational steps of the ccD-CMD algorithm to order cells along the cell cycle pseudotime (n = 10,000 cells from a HER2+ breast cancer tissue sample). (B) Histograms of the fluorescence signal of cell cycle markers measured at single-cell level. (C) Hierarchical clustering of pairwise cell cycle Difference (ccD). (D) Plot of cell cycle Difference with classical multidimensional scaling to two dimensions (ccD-CMD, left, n = 10,000 cells) with expression of cell cycle markers mapped to color (right). (E) Schematic of best-fit circle of ccD-CMD scatter (red dashed line, top) and derivation of coherence metric Inter-Octile Variation (IOV) and Circle Fit Distance (CFD). (F) Time ordering of cell cycle from ccD-CMD from G1 start (approximate G1 cell cycle start inferred from marker expression). Heat map and time plot of single-cell signal intensity measurements of cell cycle markers from the time-ordered cells (n = 10,000 cells; normalized log2 fluorescent arbitrary units, moving mean with 200 cells window). (G) ccD-CMD plot of single-cell multiplexed data of cell cycle markers from plate-based CyCIF from fixed untreated MCF10A cells (Control), serum starved for 48 hours (Starv), and exposed for 24 and 48 hours to nocodazole (Noco) or palbociclib (Palbo). Colors represent 3 distinct biological replicates, n = 1000 cells each. (H-I) Bar graph of (H) Inter-octile Variation IOV and (J) mean +/- 25^th^ percentile Circle Fit Distance CFD (KS p-value). (J) Scatterplot of cell cycle coherence metrics Inter-Octile angular Variation (IOV) and Circle Fit Distance (CFD) plots for each treatment (lines connect mean treatment values).

The molecular events driving the cell cycle are interconnected at the protein level such that fluctuations in cell cycle regulators are coordinated, and therefore correlated or anti-correlated in patterns characteristic of each cell cycle stage. To study this correlation structure we focused on epithelial cells that scored MPI +1 (i.e., proliferative tumor cells) and calculated a pairwise cell-cell correlation distance matrix in cell cycle marker space (the “cell cycle Difference” matrix; ccD; **Figure 3C**). We then transformed the data using classical multidimensional scaling (CMD) to enable visualization of the ccD in two dimensions (see Methods and Supplemental Experimental Procedures for technical details). In the resulting “ccD-CMD’’ representation, proliferating cells from a breast tumor sample formed a structure resembling a torus (**Figure 3D**). Mapping single markers on the ccD-CMD representation showed that the toroidal topography of ccD-CMD space was driven by fluctuations in the expression of cell cycle regulators (**Figure 3D**).

As a test of this approach for studying cell cycle dynamics, we applied the ccD-CMD algorithm to data generated *in silico* with a mathematical model of the cell cycle (**Figures S3E-S3H**). We used a previously described dynamical model of the mammalian cell cycle (Csikász-Nagy et al., 2006) to generate synthetic time series data, and added uncorrelated noise to simulate errors introduced during imaging of tissues. The time dynamics reconstruction by the ccD-CMD algorithm were found to be accurate, with 93% of cells being placed within 1% of their correct ordering along a canonical cell cycle trajectory (**Figure S3H**). When we compared the performance of ccD-CMD time inference with other published time inference algorithms (Cannoodt et al., 2016; Gut et al., 2015; Liang et al., 2020; Setty et al., 2019) on synthetic data and on real data from multiplexed imaging of a breast cancer cell line we found that the ccD-CMD algorithm outperformed the other inference algorithms in all settings (**Figures S3H** and **S4A-S4C**; see **Notes S5 and S6** for details on algorithm testing and comparison).

The organization of the ccD-CMD space is determined by fluctuations in the levels of cell cycle regulators, which we assume can be described by a deterministic dynamical system. If we additionally assume that our sampling is sufficient to apply the ergodic principle, we can conclude that the distance between two cells in ccD-CMD space is proportional to their difference in cell cycle position (as derived from the differential expression of the measured cell cycle regulators). A circular trajectory through the toroidal landscape of the ccD-CMD (**Figure 3D**) therefore corresponds to a prototypical progression from G1 to S to M and then back to G1. To parametrize the accuracy of this correspondence, we fit a circle to the data and derived two parameters: the uniformity of the distribution along the circumference of the ccD-CMD landscape (the inter-octile variation - IOV - in the angle θ) and the average distance of data points from the best-fit circle (circle fit distance, CFD) (**Figure 3E**). The IOV is the coefficient of variation of cell distribution in each pi/4 section of the circle. Hence, a low IOV indicates an even distribution of cells in the cell cycle. The CFD measures the dispersion of cells in ccD-CMD space: when the value is low, data from individual cells lie on or close to the best-fit circle. An even distribution of cells in ccD-CMD space is typical of unrestrained cell proliferation and corresponds to low IOV and low CFD values; we refer to this state as “cell cycle coherent” (**Figure 3E**). In coherent populations of cells, individual cells are ordered along a circle trajectory in the ccD-CMD landscape in a manner consistent with current understanding of cell cycle dynamics (Gookin et al., 2017) (**Figures 3F** and **S3I-S3K**).

To confirm our interpretation of IOV and CFD we characterized non-transformed MCF10A mammary epithelial cells grown in culture and exposed for 24 or 48 hours to the CDK4/6 inhibitor palbociclib, the microtubule inhibitor nocodazole, or serum starvation (**Figures 3G** and **S3I-S3K**). Palbociclib is expected to cause G1/S arrest, nocodazole to cause G2/M arrest and serum starvation to drive cells into quiescence. Control, untreated MCF10a cells were found to have a IOV^low^ CFD^low^ (coherent) state (**Figures 3G-3J**), the expected temporal order of cell cycle events (**Figures S3I-S3K**). Data from cells treated with palbociclib or nocodazole was skewed toward specific quadrants of the ccD-CMD landscape, representing IOV^high^ CFD^low^ states (**Figures 3G-3J**). In contrast, when cells were serum starved, data fell in a point cloud corresponding to an incoherent IOV^high^ CFD^high^ state. Thus, the higher the value of IOV and CFD the greater the deviation from freely cycling coherent proliferation (**Figures 3H-3J**).

We applied coherence metrics to cell cycle dynamics in breast cancer tissues overexpressing the HER2 growth factor receptor (Moasser, 2007). HER2 expression defines one of the major subclasses of breast cancer and is routinely assessed using immunohistochemistry (Wolff et al., 2018) as both the magnitude and heterogeneity of HER2 protein expression predict response to HER2-directed therapy (Slamon et al., 1987; Moasser, 2007; Filho et al., 2019; Katayama et al., 2021). We performed CyCIF on a cohort of 26 breast cancer specimens (TMA1, TMA2), identified MPI +1 epithelial cells and used the ccD-CMD algorithm to quantify IOV and CFD for individual patients (**Figures 4A** and **4B**). A subset of tumors were found to be in an IOV^low^ CFD^low^ state (e.g., Sample 1 in **Figure 4C**) and inspection of time-ordered cell cycle markers confirmed prototypical “coherent” cell cycle dynamics with a balanced distribution of proliferative tumor cells in both G1 and G2 phases (**Figures 4C-4E**). However, other samples deviated substantially from this pattern such as IOV^high^ Sample 2 in which cells were ‘skewed’ towards G1 and CFD^high^ Sample 3 (**Figures 4C** and **4D**). In Sample 3, cells expressed combinations of cell cycle proteins that are not found in normally cycling cells (**Figure 4E**) such as CDT1 and Geminin without detectable Ki-67, or high p21, Cyclin A, Cyclin D and phospho-Rb. Across all specimens we observed a continuum of IOV and CFD values and, within a single specimen, values also varied with gradual transitions occurring over length scales of several millimeters (**Figures 4F** and **S4G-S4K**). We combined the proliferative cells from four regions of interest (ROIs) within a single tissue and used the ccD-CMD algorithm to order cells by their cell cycle positions, thereby obtaining insight into relative cell cycle distributions in each ROI (**Figure 4G**). In ROI1, which had the highest coherence, cells were evenly distributed through the cell cycle whereas in ROI2-4, IOV was higher and cells were concentrated in different parts of the cell cycle (**Figure 4G**). Thus, proliferating breast cancers expressing the same oncogenic driver (HER2) can exhibit different cell cycle dynamics (**Figure 4A**) within a single specimen ranging from a canonical: IOV^low^ CFD^low^ state to skewed distributions consistent with cell cycle phase disruption (skewed: IOV^high^ CFD^low^) and states not normally encountered in normally growing cells (non-canonical: IOV^low^ CFD^high^).

**Figure 4.**
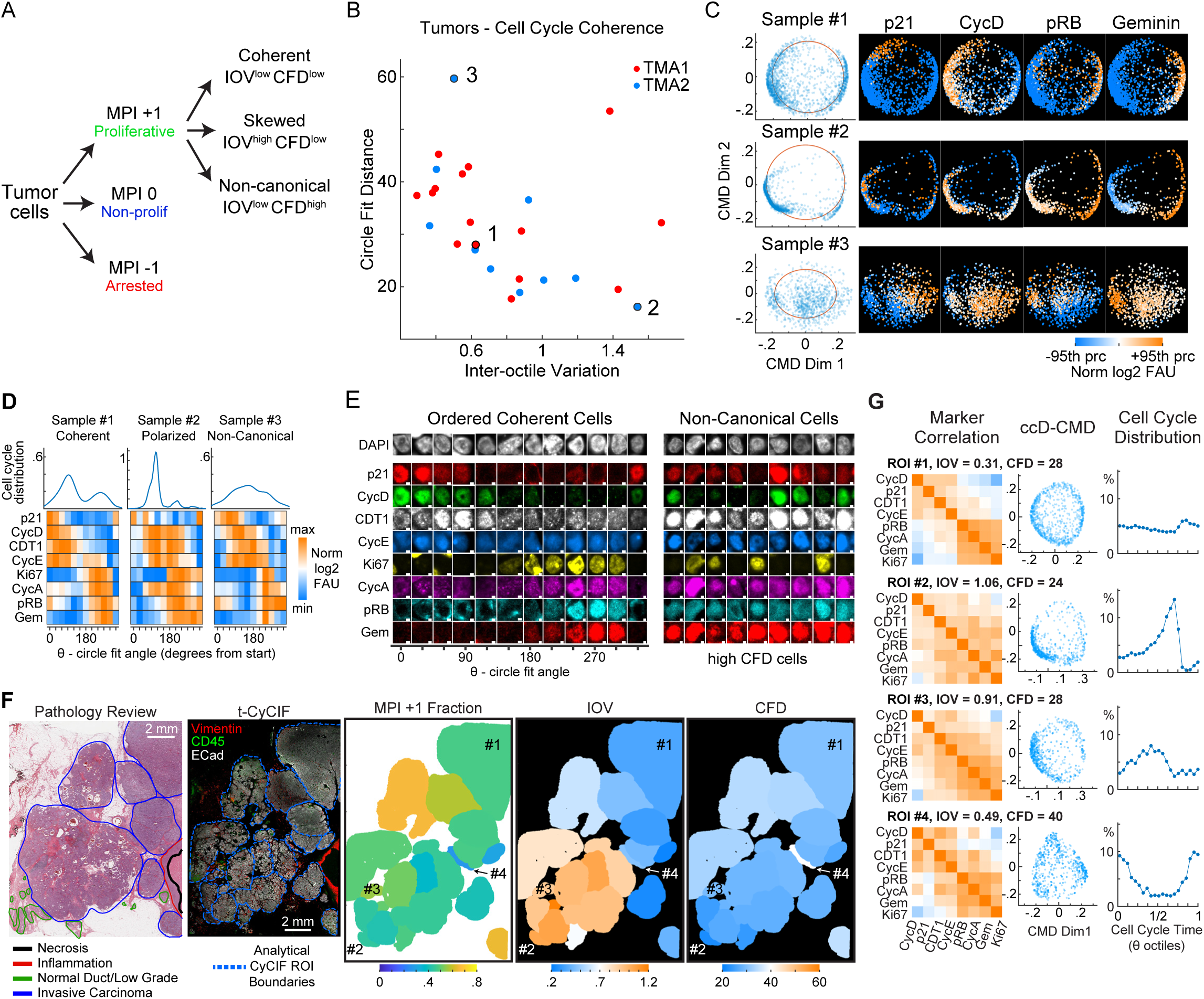
Extraction of cell cycle coherence metrics from multiplexed images of human cancer tissues. (A) Scheme to classify tumor cell populations using the MPI and characterize cell cycle coherence states. (B) Scatterplot of cell cycle coherence metrics IOV and CFD for 25 HER2+ breast samples. (C) Examples of ccD-CMD plots for the 3 indicated samples with cell cycle markers (+/-95th percentile range in sample, log2 normalized FAU). (D) Distribution of cell density and heat maps of cell cycle markers with cells binned by the circle fit angle for the 3 samples in panel C. (color, median bin intensity, normalized maximum to minimum) (E) Representative immunofluorescence images of individual tumor cells from Sample 1 (Ordered Coherent Cells) and Sample 3 (Non-Canonical Cells). Scale bar 2 µm. (F) Scanned image of hematoxylin and eosin (H&E) stained section from a HER2+ positive breast tumor with pathology annotations. Composite CyCIF images of fluorescence of tissues with annotated regions of interest (ROI) used for analysis. Spatial ROI maps of MPI and coherence metrics IOV and CFD. (G) Comparison of selected ROIs from panel F. Left, Pearson correlation matrix of cell cycle markers. Middle, cell cycle Difference and classical multidimensional scaling (ccD-CMD) plot per individual ROI. Right, plot of distribution of cells along cell cycle time from ccD-CMD performed on data from the 4 combined ROIs.

HER2 overexpression promotes proliferation of mammary epithelial cells via receptor-mediated mitogenic signaling (Goel et al., 2016; Moasser, 2007). In two independent cohorts of HER2-amplified breast tumors, we binned single cells by their HER2 protein level and assessed cell cycle dynamics using coherence metrics (TMA1 and TMA2, **Figure 5A**). Cells with the lowest HER2 levels were typically CFD^high^. Optimal coherence (IOV^low^, CFD^low^) was observed in cells expressing intermediate levels of HER2. In contrast, cells with the highest HER2 levels were also CFD^low^; their cell cycle dynamics were skewed to late G1 and IOV was high (IOV^high^, CFD^low^) (**Figures 5A** and **S5A**). Similar results were observed in individual biopsies from patients enrolled in a clinical trial (NCT02326974) of neoadjuvant dual HER2 therapy (n = 5, **Figures 5B** and **5C**).

**Figure 5.**
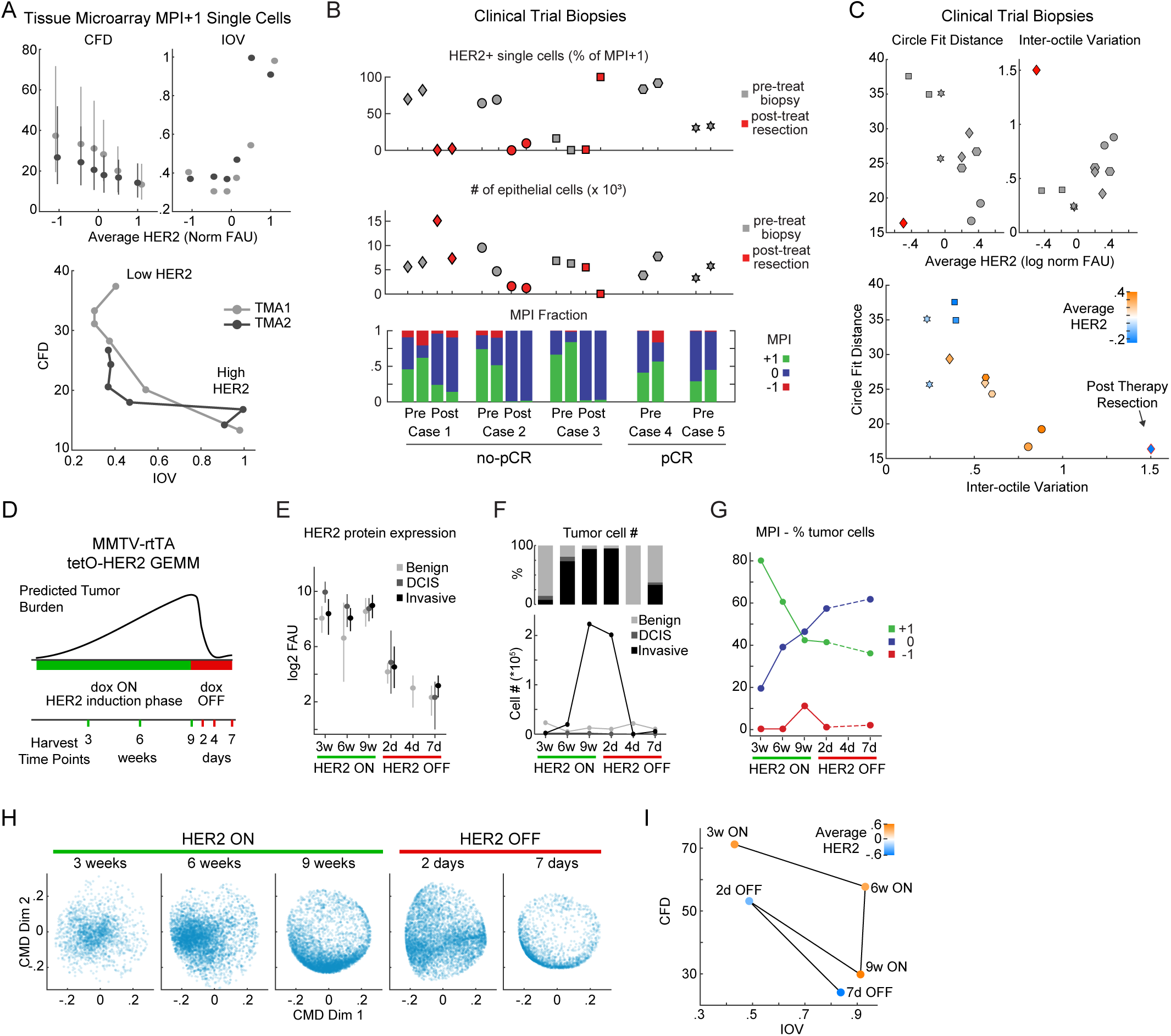
Coherence metrics are modulated by HER2 expression in human breast cancer tissues and HER2-driven mouse model of breast cancer. (A) Coherence metrics Circle Fit Distance (CFD) and Inter-Octile angular Variation (IOV) for HER2+ TMA1 and TMA2 cells (MPI +1 cells only) binned by HER2 levels (CFD mean +/- 25th percentile, n = 5000 cells per bin). Bottom, CFD versus IOV per HER2 bin, lines connect data from increasing HER2 mean levels. (B) Single-cell data summary for a 5-patient HER2+ breast cancer cohort enrolled in an anti-HER2+ clinical trial (grey markers are 2 pre-treatment biopsies, red markers are two areas of post-treatment resections). Top, percent HER2 positive cells in MPI +1 set per sample. Middle, total number of epithelial/tumor cells per sample. Bottom, fraction of cells in each MPI category. Cases 4 and 5 had a pathological complete response (pCR) to neoadjuvant therapy and did not have matched resection samples. (C) CFD and IOV versus average HER2 levels for HER2+ clinical trial samples, and plot of CFD versus IOV with HER2 level in color scale (mean log2 normalized FAU). Shapes represent patients (n = 2 biopsies per patients). Red diamond is a post-therapy resection sample. (D) Schematic diagram of HER2 induction (dox ON) and repression (dox OFF) and tissue harvest times from MMTV-rtTA tetO-HER2 genetically engineered mouse model (GEMM). (E) Plot of the median +/- 25th percentile per cell HER2 protein fluorescence in regions of normal epithelium, and in situ and invasive breast carcinoma of HER2 GEMM as defined by histology review. (F) Plot of the cell number and fraction of total cells present as benign duct epithelium, and in situ or invasive breast carcinoma in the HER2 GEMM tissues in time. (G) Plot of the fraction of tumor cells (in situ and invasive) in time in the three MPI categories. (H) ccD-CMD plot of the MPI +1 proliferative tumor cells for each time point (n = 5,000 cells per sample, day 4 did not have enough MPI +1 tumor cells). (I) CFD versus IOV for time points of the GEMM experiment.

### HER2 expression modulates cell cycle coherence in breast cancer

To determine if modulating HER2 expression alters cell cycle coherence, we used a genetically engineered mouse model (GEMM) of breast cancer in which HER2 expression can be induced (∼100-fold) and then silenced using a doxycycline-regulated breast tissue specific expression construct (MMTV-rtTA/ tetO-HER2; **Figure 5D**) (Goel et al., 2016). HER2-expressing mice develop ductal carcinoma *in situ* (DCIS) after three weeks, and palpable invasive carcinoma after a median latency of 53 days. Upon doxycycline withdrawal for 3 days (Dox-Off), tumor regression is apparent with 100% penetrance. However, more than two-thirds of mice exhibit mammary tumor recurrence within 120 days (Goel et al., 2016). As such, this model mimics important aspects of tumor dynamics in response to HER2-targeted therapy in patients, although on a faster time scale. Diverse mechanisms including upregulation of the Notch pathway, potentiation of c-Met signaling and amplification of Cyclin D1, all of which have been implicated in the development of recurrent disease in similar HER2/Neu-driven murine models (Abravanel et al., 2015; Feng et al., 2014; Goel et al., 2016).

We collected tissue samples over a nine week period of HER2 induction and seven days of subsequent HER2 repression (**Figures 5D-5G**); 27-plex CyCIF was then performed to assay tumor cell cycle dynamics. Tumors induced following nine weeks of HER2 overexpression adopted a proliferative state with a skewed cell cycle (IOV^high^ CFD^low^, **Figures 5H** and **5I**). This state resembles that of established human tumors expressing high HER2 (**Figure 5A**). Within 2 days of HER2 withdrawal, cell cycle dynamics changed to an incoherent state (IOV^low^ CFD^high^; **Figures 5H** and **5I**), even though neither proliferation (MPI +1 fraction) nor tumor cellularity had yet decreased (**Figures 5F** and **5G**). Seven days after HER2 silencing, only ∼2.5% of tumor cells remained and they exhibited skewed IOV^high^ CFD^low^ cell cycle dynamics (**Figures 5H** and **5I**). In comparison, in specimens acquired from patients enrolled in a clinical trial (NCT02326974) of neoadjuvant dual HER2 therapy, only one sample had a detectable population of proliferating cells following treatment (**Figure 5B**). In this specimen, HER2 levels were lower than in any pre-treatment sample and the state was IOV^high^ CFD^low^ (**Figure 5C**). These data suggest that the relationship between HER2 levels and cell cycle coherence is bell shaped, with the highest coherence observed at intermediate receptor levels. In both humans and mice, HER2-independent residual disease adopted the skewed cell cycle dynamics observed in pre-treatment tissues having high HER2 expression.

### Coherence metrics change with treatment and are associated with clinical outcome

As an additional means of studying how therapy impacts coherence, we assayed specimens from three patients with localized breast cancer who were biopsied before, during and after treatment (patient samples also analyzed in **Figure 2E**). In one patient with triple negative breast cancer (TNBC) biopsied prior to treatment (pre), following 12 weeks of neoadjuvant paclitaxel (on), and then again after 20 additional weeks of treatment with doxorubicin-cyclophosphamide (post) (**Figure 6A**), we found that either paclitaxel or doxorubicin-cyclophosphamide induced only small changes in the MPI +1 fraction (**Figure 6B**), even though the Ki-67+ fraction fell ∼50% in the on-paclitaxel specimen (**Figure 6C**). ccD-CMD analysis of the three longitudinal TNBC samples showed that the cell cycle dynamics were mostly coherent throughout the treatment (CFD < 40, IOV < 0.6, **Figure 6D**). However, the dynamics of the “on” sample skewed towards the G1 phase of the cell cycle (**Figures 6E-6G**), consistent with data from intravital imaging of xenograft models treated with paclitaxel (Chittajallu et al., 2015). G1 accumulation in the presence of paclitaxel explains why Ki-67 staining alone underestimated the proliferative fraction (**Figure 6C**). In another type of breast cancer (ER^+^ cancer) two other “pre-on-post” sample sets collected longitudinally over time showed drastic changes in coherence metrics induced by therapy (**Figures 6H** and **6I**), even when treatment lasted for as little as two weeks (“pre” to “on” samples). Changes in coherence were independent of changes in the fraction of proliferating cells (MPI +1 fraction; **Figure 6H**). We conclude that cell cycle coherence is a plastic phenotype that provides a sensitive measure of therapy-induced changes independent of significant reductions in proliferative index.

**Figure 6.**
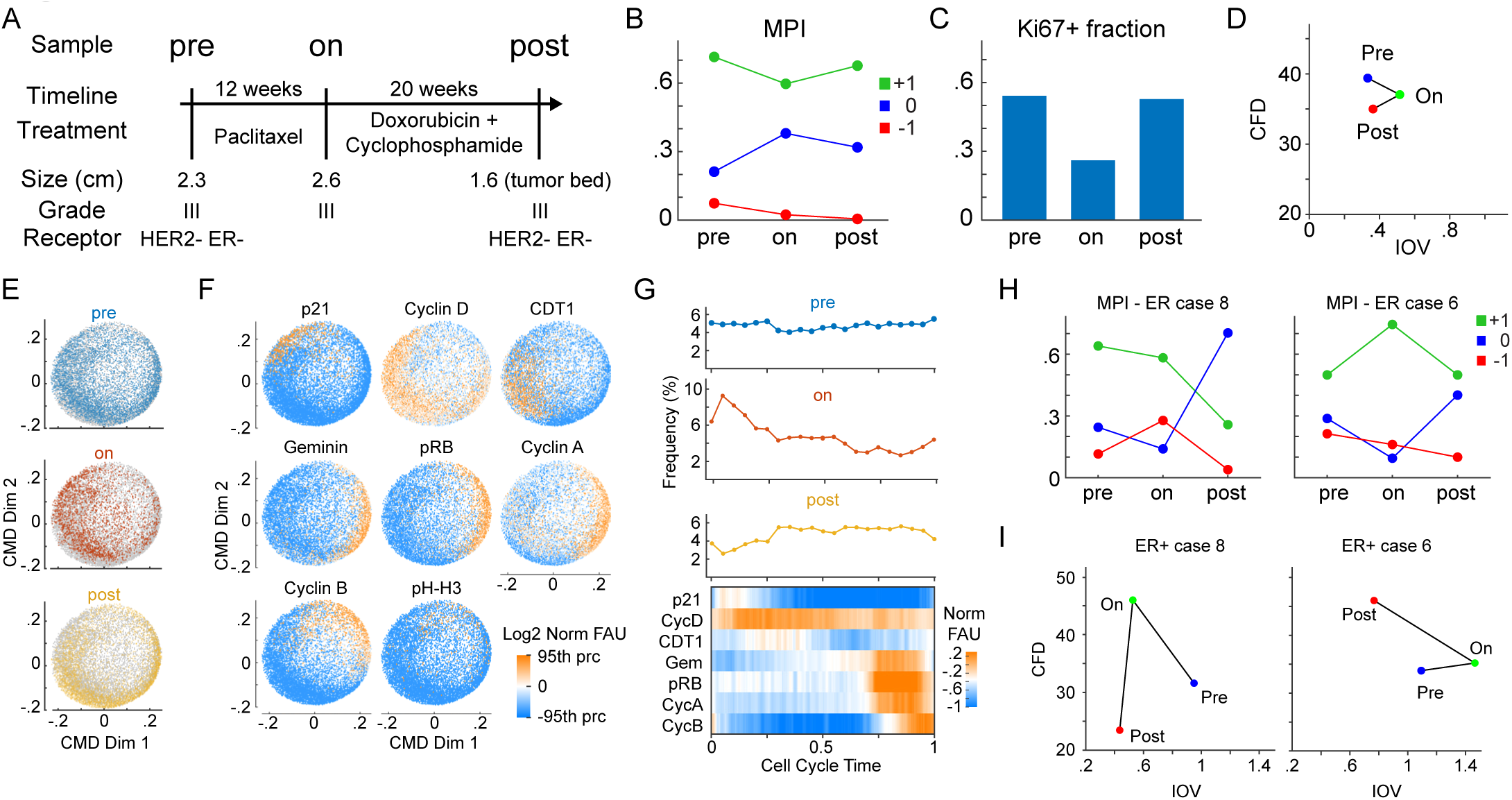
Coherence metrics change with treatment. (A) Clinical and pathologic features of biopsy/resection tissues from a patient with triple negative breast cancer (TNBC) characterized by CyCIF imaging. Three samples include the diagnostic biopsy (pre) and two samples after indicated treatments (on, post). (B-C) Plot of (B) fraction of cells in the three MPI categories and (C) Ki-67 positive fraction, through the treatment course for samples in panel A. (D) Scatterplot of coherence metrics Circle Fit Distance (CFD) versus Inter-Octile angular Variation (IOV) for triple-negative breast cancer (TNBC) patient (pre n = 5,000, on n=2,250, post n = 5,000 single MPI +1 cells). (E-F) ccD-CMD plot of combined data from TNBC pre, on, and post samples with data (E) corresponding to time of biopsy indicated by color, and (F) with single marker normalized intensities mapped to color (n = 5,000 cells per plot, pH-H3 was not used by the ccD-CMD algorithm). (G) Distribution of cells along cell cycle time in the three samples and heat map of marker expression for single cells across cell cycle time combined for the three samples (moving mean over 100 cells). Upper panels, single time point cell frequency distribution. (H) Plot of fraction of cells in the three MPI categories through the treatment course for samples from two ER+ breast cancer patients pre-, on-, and post-treatment (see **Table S5** for treatment details). (I) Scatterplot of coherence metrics Circle Fit Distance (CFD) versus Inter-Octile angular Variation (IOV) for two ER+ breast cancer patients pre-, on-, and post-treatment from panel H.

To determine if differences in cell cycle coherence are associated with differences in disease outcome we assayed specimens from patients cohorts diagnosed with two different lethal malignancies (mesothelioma and glioblastoma). Patients were stratified into a coherent IOV^low^, CFD^low^ group and an incoherent group encompassing either IOV^high^ or CFD^high^ states (**Figures 7A** and **7B**). We found that patients whose tumors exhibited coherent cell cycle dynamics had significantly worse outcomes (logrank p-value <0.02). Similar results were obtained if tumors were stratified into three groups (coherent, skewed and non-canonical; **Figure S6**). We conclude that cycle coherence in mesothelioma and glioblastoma is associated with aggressive tumor behavior and worse progression free survival.

**Figure 7.**
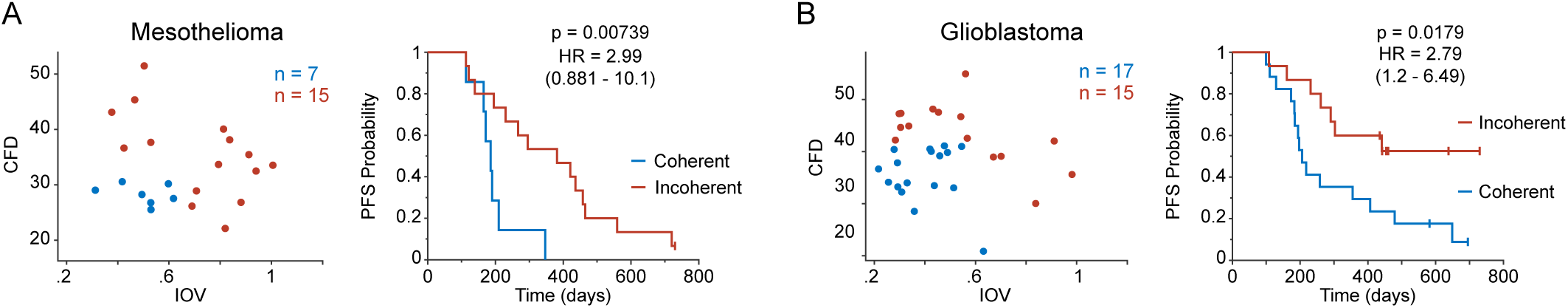
Coherence metrics are associated with clinical outcome. (A) Left, scatterplot of CFD versus IOV from a mesothelioma sample cohort (n = 22 patients). Colors represent binning into coherence groups according IOV and CFD metrics. Right, corresponding Kaplan Meier estimation of progression-free survival (PFS) for the patients (logrank p-value). (B) Left, scatterplot of CFD versus IOV from a glioblastoma sample cohort (n = 32 patients). Colors represent binning into coherence groups according IOV and CFD metrics. Right, corresponding Kaplan Meier estimation of progression-free survival (PFS) for the patients (logrank p-value).

## DISCUSSION

Experimental models of cancer have shown that cell cycle progression and tumor cell proliferation are dynamically regulated processes, influenced by both cell autonomous and non-autonomous factors (Mahdessian et al., 2021; Hanahan and Weinberg, 2011, 2000). These properties are largely unexplored in the setting of intact human tissue and tumors. Instead, proliferative fraction is usually captured via a dichotomous assessment of the levels of a single marker (Ki-67). This approach incorrectly makes some dividing cells appear non-proliferative and fails to capture the cell cycle heterogeneity of proliferating cells. Here we provide a more informative and potentially more robust molecular characterization of proliferation that lays a foundation for understanding how tumors integrate microenvironmental and cell-intrinsic signals to grow in different niches.

Specifically, a multivariate proliferation index (MPI) provides a means to quantity proliferative index based on the expression of multiple marker proteins – not just Ki-67 – and cell cycle coherence measures how closely cell cycle dynamics conform to those of the well understood cell cycles in freely dividing cancer cells. Through the lens of coherence, we show that only a subset of tumors grow with ”canonical” dynamics while others are skewed towards specific cell cycle phases or express unanticipated combinations of regulators. In both humans and mice, we find that intermediate levels of HER2 expression promote coherent cell cycle dynamics, whereas higher levels of oncogene overexpression lead to skewed dynamics, potentially resulting from phase-specific deceleration or acceleration involving skipped restriction points (Min et al., 2020; Moser et al., 2018). High coherence may not necessarily translate into faster growth, but both skewed and non-canonical dynamics are associated with less aggressive tumors in two highly lethal malignancies.

We observe diverse cell cycle states within and across specimens. By contrast, the spatial organization of cells into proliferative and non-proliferative domains of two characteristic lengths appears to be conserved across cancers of diverse histology. We speculate that small-scale proliferative structures correspond to cells and their daughters following a few divisions (and perhaps also mitogenic cell-cell interactions) whereas large neighborhoods arise from differences in environmental conditions and structural constraints. These neighborhoods contain thousands of either proliferating or non-proliferating cells, reminiscent of developmental patterning by morphogen gradients (Briscoe and Small, 2015).

Basic research, clinical trials, and precision cancer medicine require new quantitative measurements and computational approaches to correctly quantify cancer cell states and phenotypes, and their spatial organization in tissues (HuBMAP Consortium, 2019; Rajewsky et al., 2020; Rozenblatt-Rosen et al., 2020). This information is orthogonal but complementary to the characterization of genomic heterogeneity in tumors, and is expected to provide new means to understand response to treatment and the evolution of drug resistance (Caswell-Jin et al., 2019; Lomakin et al., 2021; Rueda et al., 2019; Zahir et al., 2020).

## ACKNOWLEDGEMENTS

This work was supported by NIH grants R01-CA194005 (SS), R41-CA224503 (PKS), U54-CA225088 (PKS, SS), P50-CA168504 (J.J.Z) and R35-CA210057(J.J.Z.); DF/HCC Breast SPORE: Specialized Program of Research Excellence (SPORE), P50-CA168504 (EPW); Ludwig Center at Harvard (PKS, SS), T32-HL007627 (GG), and the American-Italian Cancer Foundation postdoctoral fellowship (GG); Terri Brodeur Breast Cancer Foundation (SK), the Breast Cancer Research Foundation BCRF (J.J.Z.), DOD CDMRP W81XWH-18-1-0491 (J.J.Z.). We thank Dana-Farber/Harvard Cancer Center for the use of the Specialized Histopathology Core, which provided histopathology services. Dana-Farber/Harvard Cancer Center is supported in part by an NCI Cancer Center Support Grant P30-CA06516. We thank Clarence Yapp and Yu-An Chen for assistance with microscopy and image analysis, and Chris Rycroft and Galit Lahav for helpful discussions.

## AUTHOR CONTRIBUTIONS

Conceptualization – GG, SK, JJZ, PKS, SS

Methodology – GG, SK, JB, SC, JRL, PKS, SS

Software – GG, DA, YD

Validation – GG, SK, DA, YD, SC, DAD

Formal Analysis – GG, SK, DA, YD, SC

Investigation – GG, SK, JJZ, PKS, SS

Resources – EPW, JJZ, PKS, SS

Data Curation – GG, SK, DA, YD, SC, DAD

Writing – Original Draft – GG, SK, SS

Writing – Reviewing & Editing – GG, SK, DA, YD, JJZ, PKS, SS

Visualization – GG, SK, YD

Supervision – JJZ, PKS, SS

Project Administration – PKS, SS

Funding Acquisition – EPW, JJZ, PKS, SS

## METHODS

### RESOURCE AVAILABILITY

#### Lead contact

Further information and requests for resources and reagents should be directed to and will be fulfilled by the lead contacts, Sandro Santagata and Sheheryar Kabraji.

#### Materials availability

This study did not generate any unique reagents.

#### Data and code availability

Datasets generated and the corresponding analysis used in all figures are available upon request.

### EXPERIMENTAL MODEL AND SUBJECT DETAILS

#### Cell Lines

MCF10A (female mammary epithelial) cells were purchased from ATCC and grown in DMEM/F12 Medium (Gibco, 11330-032) with 5% horse serum (Gibco, 16050-122), 1% penicillin/streptomycin (Gibco, 15070-063), 0.1% Insulin (Sigma, #I1882), 0.05% hydrocortisone (Sigma, #H0888), 0.02% human-EGF (Sigma, E5036), and 0.01% cholera toxin (Sigma, #C8052). Cells were incubated at 37°C and 5% CO2 until fixation.

#### Human Tissue Sections

Whole slide tissue sections of breast carcinoma, ovarian carcinoma, and squamous cell lung carcinoma, and sections of tissue microarrays (TMAs) of breast carcinoma (HTMA 226, 227; triplicate mm diameter cores per case; courtesy of the DFCI Breast Oncology Group), glioma (HTMA 399; quadruplicate 0.6 mm cores per case), colorectal carcinoma (HTMA402; triplicate 0.6 mm cores per case), and mesothelioma (HTMA403, triplicate 1.0 mm cores per case) were prepared from formalin fixed, paraffin embedded (FFPE) tissue blocks from sample retrieved from archives of the Department of Pathology at Brigham and Women’s Hospital (BWH) in accordance with Institutional Review Board (IRB) approved protocols at BWH and the Dana Farber Cancer Institute. See **Table S3** for sample clinical information of cases analyzed as whole tissue sections. A outcome analysis of progression free survival was performed on a subset of the cases retrospectively collected from HTMA399 and HTMA403 and included only patients with clinical follow up following resection of primary tumors that were treated using standard of care regimens (therapy information in **Tables S6-S7**) and in the case of gliomas (HTMA399), only IDH wild type glioblastoma were included in the analysis. FFPE tissue sections of breast carcinoma tissue microarray BRC15010 were purchased from Pantomics, Inc. Clinical data such as gender, age, therapy, and diagnosis for the over 700 patients whose samples are used in this paper can be found in the **Tables S8-S10**.

#### Animal experiments

MMTV-rtTA/tetO-HER2 mice were previously generated (Goel et al., 2016). The transgenic construct was induced by introducing a doxycycline containing diet to 8 week-old female FVB MMTV-rtTA/tetO-HER2 mice as previously described (Goel et al., 2016). Two mice were sacrificed at 3, 6, and 9 weeks following introduction of the doxycycline diet and 2, 4, and 7 days after withdrawal of the doxycycline by switching to a standard diet. Mice were euthanized using CO2 inhalation, and all mouse experiments were performed in accordance with protocol 06-034 approved by the Institutional Animal Care and Use Committees of Dana-Farber Cancer Institute and Harvard Medical School. Multiple primary tumors were excised from each mouse and processed into FFPE tissue blocks.

## METHOD DETAILS

### Experimental methods

#### Plate-based Cyclic Immunofluorescence (p-CyCIF)

MCF10A cells were grown and plated in flat bottom polystyrene 96-well plates at approximately 1E6 cells/mL. Two plates were seeded at once for the two treatment times, and 12 wells were seeded to account for triplicates of each of 4 treatments. After 24 hours, cells were either given fresh media or treated with serum-free media, 1 µM palbociclib (Sigma, #PZ0383), or 5 µM nocodazole (Cell Signaling Technology, 2190S) and incubated at 37°C for 24 or 48 hours. After treatment, Click-iT™ EdU Alexa Fluor™ 488 solution (Molecular Probes, PZ0383) was added to all the wells to a final concentration of 10 µM and incubated at 37°C for 2 hours. The cells were washed in Dulbecco’s phosphate-buffered saline (DPBS; Gibco, 14190-250) and fixed with 3.7% paraformaldehyde (Electron Microscopy Science, C993M23) for 30 minutes. Cells were washed and permeabilized with 0.5% Triton® X-100 (Sigma, X100) in PBS (Gibco, 10010023) for 15 minutes.

Plate-based Cyclic Immunofluorescence (p-CyCIF) was performed as previously described in detail (Lin et al., 2015). Briefly, fixed wells underwent multiple cycles of incubation with primary-labeled antibody, imaging, and fluorophore inactivation. Three primary conjugated antibodies and Hoechst 33342 (Thermo Fisher Scientific, 62249) were diluted in Odyssey Blocking Buffer (LI-Cor, cat. no. P/N 927–40003) and incubated at 4°C overnight protected from light. See **Table S4** for a complete list of antibodies used in each experiment and their dilutions; antibody dilutions range from 1:10 to 1:500 with 1:100 being the most common. Cells were washed and placed in PBS for imaging using DAPI, FITC, Cy3, and Cy5 channels on a GE IN Cell Analyzer 6000. The fluorophores were inactivated with 200 µL of bleaching solution (4.5% H2O2 and 20 mM NaOH in PBS; Sigma, 216763, Sigma, 221465) for 1 hour under LED lights (Amazon, cat. no. B078JCBW9S) and washed before a new cycle of antibodies was added. After completing all rounds of p-CyCIF, the EdU Click-iT Reaction (Molecular Probes, PZ0383) was performed according to the manufacturer’s protocol and imaged as the previous rounds.

#### Tissue Cyclic Immunofluorescence (t-CyCIF)

Tissue Cyclic Immunofluorescence (t-CyCIF) was conducted as previously described (Du et al., 2019; Lin et al., 2018) on the full slide tissues and tissue microarrays. In preparation for t-CyCIF, the FFPE slides were baked at 60°C for 30 minutes, dewaxed using Bond Dewax solution at 72°C, and antigen retrieval was performed with Epitope Retrieval 1 solution at 100°C for 20 minutes using the BOND RX Automated IHC/ISH Stainer. Each round’s antibodies were diluted in Odyssey Blocking Buffer and incubated overnight at 4°C in the dark. See **Table S4** for the complete list of antibodies. After antibody incubation, slides were stained with Hoechst 33342 for 10 minutes at room temperature. Slides were coverslipped using 20-50% glycerol solution (Sigma, G5516) in PBS. Images were taken using DAPI, FITC, Cy3, and Cy5 channels either on the GE IN Cell Analyzer 6000 (20x/0.75NA objective lens) or on the RareCyte CyteFinder (20x/0.75NA objective lens). After imaging, the fluorophores were inactivated with bleaching solution (4.5% H2O2 and 20 mM NaOH in PBS) for 45 minutes under LED lights, and the cycle was repeated.

### Image processing

The image processing of both plate-based and tissue cyclic immunofluorescence is organized in the following steps, each of which is described in detail below:

i. the software ASHLAR is used to stitch, register, and correct for image acquisition artifacts (using the BaSiC algorithm). The output of ASHLAR is a single pyramid ome.tiff file for each region imaged;
ii. the ome.tiff file is re-cut into tiles (typically 5000 x 5000 pixels) containing only the highest resolution image for all channels. One random cropped image (250 x 250 pixels) per tile is outputted for segmentation training (using Fiji);

a. note that for plate-based experiments that steps i-ii) are optional
iii. the ilastik software is trained on the cropped images to label, nuclear, cytoplasmic, and background areas. The output of the Ilastik processing is a 3-color RGB image with label probabilities;
iv. the RBG probability images are thresholded and watershed in MATLAB to segment the nuclear area. The cytoplasmic measurements are derived by dilating the nuclear mask;
v. single-cell measurements are extracted for each channel (cell pixel median and mean for both nuclear and cytoplasmic area) as well as morphological measurements of area, solidity, and cell coordinates location.

#### BaSiC

The BaSiC ImageJ plugin tool was used to perform background and shading correction of the original images (Peng et al., 2017). The BaSiC algorithm calculates the flatfield, the change in effective illumination across an image, and the darkfield, which captures the camera offset and thermal noise. The dark field correction image is subtracted from the original image, and the result is divided by the flatfield image correction to obtain the final image.

#### ASHLAR

Alignment by Simultaneous Harmonization of Layer/Adjacency Registration (ASHLAR) is used to stitch together image tiles and register image tiles in subsequent layers to those in the first layer (Muhlich et al., 2021). For the first image layer, neighboring image tiles are aligned to one another via a phase correlation algorithm that corrected for local state positioning error. A similar method is applied for subsequent layers to align tiles to their corresponding tile in the first layer. ASHLAR outputs an OME-TIFF file containing a multi-channel mosaic of the full image across all imaging cycles. Full codes available at: https://github.com/labsyspharm/ashlar.

#### ilastik

ilastik is a machine learning based bioimage analysis tool that is used to obtain nuclear and cytoplasmic segmentation masks from OME-TIFF files (Berg et al., 2019). For increased processing speed, randomly selected 250 x 250 pixel regions from the original OME-TIFF are used as training data. ilastik’s interactive user interface allows the user to provide training annotations on the cropped regions. Users are presented with a subset of the channels stacked images and label pixels as either nuclear area, cytoplasmic area, or background area. The annotations are used to train non-linear classifiers that are applied to the entire image to obtain probability masks describing the probabilities of each pixel belonging to the nuclear, cytoplasmic, or background area. A MATLAB (version 2018a) script uses these masks to construct binary masks for nuclear and cytoplasmic area.

### Data analysis workflow

The data analysis is divided in a set of pre-processing steps in which data from different tissues is i) log2-transformed and aggregated together, ii) filtered for image analysis errors, and iii) normalized on a channel-by-channel basis across the entire data from a single experiment. All the steps are performed in MATLAB.

#### Data aggregation

The image processing workflow outputs one ome.tiff image and one data file (.mat) for each tissue area imaged. The data matrices from each .mat file are concatenated into a single matrix for each metric measured (median/mean, nuclear/cytoplasmic) into a single structure (“AggrResults”). The morphological data (i.e., area, solidity, and centroid coordinates) is concatenated into a single structure (“MorpResults”), which also contains the indexing vector to keep track of the tissue of origin within the dataset.

#### Data filtering

Single cells are filtered to identify and potentially exclude from subsequent analysis errors in segmentation and cells lost through the rounds of imaging. Two types of criteria are used to filter cells: morphological criteria based on cell object segmented area, which are applied to all the rounds for the cell object, and DAPI-based criteria which are applied to the DAPI measurement for each imaging round. The latter corrects for cell loss during cycling and computational misalignment, which are both round specific.

Morphological filtering criteria are:

- nuclear area within a user-input range;
- cytoplasmic area within a user-input range;
- nuclear object solidity above a user-input threshold.

DAPI-based criteria are:

- nuclear DAPI measurement above a user-input threshold;
- ratio between nuclear and cytoplasmic DAPI measurement above a user-input threshold; The filter information for the criteria is allocated to a logical (0-1) structure ‘Filter’, which is used to select the cells to analyze in the further analysis by indexing. The threshold selection is dataset dependent and is performed by data inspection. The values used in each dataset are available upon request.

#### Data normalization

Each channel distribution is normalized by probability density function (pdf) centering and rescaling. The aim is to center the distribution of the log2 fluorescent signal at 0 and rescale the width of the distribution to be able to compare across channels. The data is first log-transformed (base 2). The standard normalization is performed using a 2-component Gaussian mixture model, each model capturing the negative and the positive cell population. If the 2-component model fails to approximate the channel distribution, two other strategies are attempted: i) a 3-component model is used assuming the components with the two highest means are the negative and positive distribution (i.e., discarding the lowest component) or ii) the user selects a percentage ‘x’ of assumed positive cells and a single Gaussian distribution fit is performed on the remainder of the data to capture the negative distribution. The single Gaussian fit is then used as the lower component in a 2-component model to estimate the distribution of the positive population. The “add_coeff” is defined as the intersection of the negative and positive distributions. The “mult_coeff” is defined as the difference between the mean of the negative and positive distributions. The full distribution is normalized by subtracting the add_coeff and dividing by the mult_coeff. The normalization is performed on the nuclear and cytoplasmic single-cell, single-channel distributions individually.

The data preprocessing workflow is performed on all datasets. The individual analyses used in the paper are performed only in selected datasets as follows.

#### Cell type calling strategy

Cells from tissue based experiments are separated into lineage compartment by cell type markers. Cells are scored based on the sign of the normalized value using the following criteria:

- epithelial cells, positive for e-cadherin OR pan-cytokeratin;
- immune cells, positive for one of CD45, CD3D, CD4, CD68, CD163, CD8a;
- stromal cells, positive for αSMA and negative for epithelial markers OR positive for Vimentin and negative for immune ^α^markers;
- others/not classifiable, negative for all the markers in the categories above

Note that not all markers were imaged in all datasets, hence only the available ones were used. If a cell was called as more than one single cell-type, this is defined as a conflict. The conflicts are resolved by comparing the markers that triggered each of the cell type calls and assign the cell type with the highest marker level. If the markers are within 10% of each other the cell is deemed “not classifiable”.

#### Multivariate proliferative index (MPI) calculation

The multivariate proliferation index, or MPI, is a based on the normalized measurement of 5 markers that are used in all the tissue-derived datasets in this work: three markers of proliferation, Ki-67, MCM2, and PCNA and two markers of cell cycle arrest, p21 and p27. The aim was to capture proliferation robustly, avoiding relying on a single marker, while also separating cells expressing high levels of arrest markers (even if they express markers of proliferation). The logic for the MPI determination is:

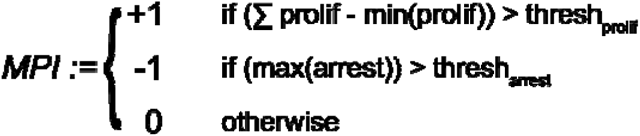

The determination of the threshold values for the proliferation and arrest is dataset dependent. However due to our marker normalization strategy the values were comparable between datasets (thresh_prolif_ = 0 and thresh_arrest_ ∼ 0.25-0.75, tuned based on Ki-67 levels).

#### Clustering and t-SNE

Clustering was performed using the k-means algorithm. The cluster arrangement was determined by hierarchical clustering of k-means clustering mean value per cluster. Both clustering were performed in MATLAB using in-built functions “kmeans” and “clustergram”. The t-distributed stochastic neighbor embedding (t-SNE) was performed on a subset of cells (specified in figure legends) with perplexity parameter set at 500.

#### Spatial Correlation Analysis

Spatial correlations C_xy_(r) were computed as the Pearson correlation between a cell of group X and its k^th^ nearest neighbor of group Y, for their respective variables x and y. A value of C_xy_ (r) was computed for each k up to 100, and a distance r was assigned to each k as the average distance between k^th^ nearest neighbors. For the MPI spatial correlation cell groups X and Y were both epithelial/tumor cells, and variables x and y were logical, whether the cell belonged to the specified MPI category or not. For the correlation between the MPI categories and the tumor microenvironment, group X was the epithelial/tumor cells and group Y was the immune compartments. The variable x and y were logical, x = MPI category, y = subtype of immune cells. To calculate the characteristic lengths l_1_ and l_2_, the C_xy_ data was fitted with a two exponential fit y = a_1_*exp(-x/l_1_) + a_2_*exp(-x/l_2_) by least-square fitting with all parameters constrained to be positive (MATLAB in-built function lsqcurvefit). The estimates derived from tissue microarrays (TMA) cores were filtered for fits with residuals below 0.05 as calculated by lsqcurvefit function. The long range estimates were also filtered for values below the core size (1.1 mm for mesothelioma cores and 600 microns for all other TMAs).

#### Upset plot

The Upset plots were computed using the R-package available at https://caleydo.org/tools/upset/, developed by the Visual Science Data, Institute of Computer Graphics at Johannes Kepler University Linz, Austria.

#### Pairwise cell cycle difference (ccD) and classical multidimensional scaling reduction (CMD)

The ccD is calculated by taking the absolute value of the pairwise Pearson correlation between the cell cycle marker vector of normalized values of each cell. The CMD was performed with a MATLAB built-in function cmdscale. The ccD was reshaped to fit the requirements expected by the cmdscale algorithm, ccD’ = 0.5-ccD/2. The first two dimensions of the CMD scaling are plotted and used for further analysis.

#### Circular fit, cell cycle dynamics reconstruction, and cell cycle coherence summary metrics

For both the simulation data and the plate-based and tissue CyCIF data the same algorithm was followed. The CMD scaled two-dimensional data is fit by least-squares minimization to a circle. For each point in the data two parameters are calculated, i) the distance to the closest point of the circle (circle fit distance, CFD) and ii) the angle of the point to the origin of the fitted circle. The angle is used to order the cells in what is referred to as the “cell cycle ordering”. Given the cyclical structure of the ordering, the origin (i.e., time 0) is arbitrary and was set to separate the M phase marker pattern from the early G1 phase pattern. The cell cycle distribution plots (**Figure S5**) are computed by aggregating cells from the ROI listed (5000 cells max for each ROI), running the ccD-CMD algorithm to order cells along the cell cycle, quantifying the histogram of cells belonging to each ROI and normalizing the frequency to the cell number of the specific ROI.

The cell cycle coherence distance is the circular fit distance detailed above. The angular distribution coefficient of variation is calculated by binning the angle measurements (ii) above into 8 bins and calculating the proportion of cells in each bin. The proportion calculation is repeated by shifting the bin position by π/8 to ensure lack of positional bias in the bin definition. We hence refer to this metric as the inter-octi^π^le angular variation (IOV). The IOV is the coefficient of variation of the bin proportions (which is equal to 0 in a uniformly distributed population).

A comparison between the ccD-CMD pseudotime cell cycle ordering and three previously published time inference algorithms is detailed in the **Note S6** and in **Figure S3 and S4.**

#### Outcome analysis

The outcome analysis was performed using Kaplan-Meyer estimation and logrank test. The analysis was computed in MATLAB using the MatSurv function (Creed et al., 2020). The cutoffs were chosen based on cell line observation (IOV = 0.62, CFD = 42) or using the median value for Ki-67 cutoff. The CFD threshold for the mesothelioma cohort was lowered to 37 in order for the “non-canonical” category to have a minimum of 5 patients. Analysis was restricted to cases with >500 MPI+1 cells. The treatment received is listed in **Tables S6-S7**. Only patients that received chemotherapy were included in the outcome analyses.

### Cell cycle modeling

The cell cycle was modeled *in silico* using a system of ordinary differential equations (ODEs) based on the model by Csikász-Nagy and Tyson (Csikász-Nagy et al., 2006). A Python script utilizing Euler’s method was used to solve the ODEs. Measurement noise sampled from a Poisson distribution was introduced to reproduce background from microscopy experimental settings, to include both the shot noise and measurement fluctuations. For the “untreated” conditions, the ODEs, kinetic constants, and initial values were based on published parameters (Csikász-Nagy et al., 2006), with the only variation being the lowering of the “maxmass” parameter to 1.8. The “G1 arrest” condition was simulated in the model by setting the values of the active CDK/cyclinD complexes to zero. This is used to model the molecular mechanism of the drug palbociclib.

## QUANTIFICATION AND STATISTICAL ANALYSIS

Information on the sample size, the number of independent repeats that were used to derive statistics, the statistics used to summarize the data for each experiment is presented in Table S1. Curve fitting was performed using MATLAB (except for linear fit in Figure S1E which was performed in MS Excel). Statistical tests used are Pearson correlation, two-sided t-test, Kolmogorov–Smirnov (KS) and logrank as specified in the figure legends (**Figures 1G, 2G and 2I, 3H-3I, 7A-7B, S1F, S2D-S2G,** and **S6**) and are performed with MATLAB built-in functions. Only samples with a minimum of 100 single cells (after quality control filtering) were included in the analyses. Significance was defined as a p-value of less than 0.05.

## ADDITIONAL RESOURCES

Multiplexed images of human HER2 breast cancer used in **Figure 3A** can be viewed in *Minerva Story* (Hoffer et al., 2020; Rashid et al., 2020) an interpretive guide for interacting with multiplexed tissue imaging data https://tinyurl.com/minerva-proliferation

## SUPPLEMENTAL INFORMATION

Supplemental Information includes seven figures, ten tables, and six notes, and can be found with this article online.

**Table S1:** summary and statistics of CyCIF datasets.

**Table S2:** cell cycle markers used for each sets of data.

**Table S3:** clinical information about patient samples.

**Table S4:** list of antibodies used in the study.

**Table S5:** clinical information on biopsy samples from breast cancer patients.

**Table S6:** clinical data on glioblastoma cohort used in outcome studies.

**Table S7:** clinical data on mesothelioma cohort used in outcome studies.

**Table S8:** human samples clinical data - Demographics

**Table S9:** human samples clinical data - Diagnosis

**Table S10:** human samples clinical data – Therapy

## SUPPLEMENTAL NOTES

**Supplemental Note S1: Current tissue-based proliferation scoring metrics**

In clinical settings and research studies using tissues, cellular proliferation is typically scored by the expression of a single protein marker assayed by immunohistochemistry (IHC). The most commonly used markers of proliferation are Ki-67, phospho-RB, Cyclin D, and the “mitotic index”, which is a count of the number of mitotic figures per defined area from haematoxylin and eosin stains. Clinical scoring is often performed manually by a pathologist that visually reviews IHC images and assigns an overall proliferation score per tissue or is occasionally performed using semi-automated assessment. These approaches have a number of limitations. First, the manual scoring by pathologists is subject to human bias, and is not quantitative or scalable. Second, single markers are not able to accurately capture proliferation. The markers currently used are biased toward specific parts of the cell cycle; Ki-67 and phospho-RB preferentially stain cells in the S/G2 phase of the cell cycle, Cyclin D stains cells in G1 and the mitotic index reports for cells in M phase. These markers have a low false positive rate - they are unlikely to stain non-proliferating cells - but they are typically expressed in specific phases of the cell cycle only. Other markers of proliferation, such as PCNA and MCM2 are more ubiquitously expressed during the cell cycle and are more stable proteins. However, they are co-expressed with markers of cell cycle arrest (Figures 1A, 1B, S1B, and S1C), and hence have a higher rate of false positives. Lastly, single stains for proliferation do not allow one to discriminate between the various cell types present in the tumor areas. Both infiltrating stromal and immune cells proliferate within the tumor microenvironment and cannot be visually excluded from the scoring and may confound the measurement of cancer cell proliferation.

**Supplemental Note S2: Comparison between the MPI and other multidimensional classification strategies**

In Figures 1E and S1D we show a direct comparison between MPI calls and both k-means clustering and t-distributed stochastic neighbor embedding (t-SNE), two commonly used tools to classify multidimensional dataset from various multiplex techniques (scRNAseq, CyTOF, MIBI, IMC, CODEX). The concordance between the MPI single cell calls and k-means clustering is evident by the fact that the majority of clusters identified by the k-means algorithm consist mainly of a single MPI category. K-means clustering still involves manual selection of the number of clusters and manual definition of each cluster’s identity, which would have to be inspected for each experiment individually. Further, while the MPI definition is deterministic, clustering has a stochastic component that changes the cluster assignment every time the algorithm is run, which makes it not fully reproducible and less scalable.

The t-SNE results are also concordant with the MPI classification, with MPI cells from different categories segregating away from each other (Figure 1E, the MPI was not used in input for the t-SNE algorithm). Classifying cells on the t-SNE representation still requires running a clustering algorithm on the t-SNE output (DBSCAN is often used for this purpose), which leads to similar problems as mentioned above for k-means clustering. The major drawback of t-SNE and similar dimensionality reduction algorithms is that they are not computationally scalable and cannot be run on the large datasets generated in our study (on the order of hundreds of thousands to millions of cells). While there exist machine learning techniques that allow building classification models from subsets of the data, these were beyond the scope of this study and did not present clear advantages on the single cell MPI deterministic classification strategies.

**Supplemental Note S3: Lack of accurate quantification of total DNA and DNA replication**

A unique limitation in the study of proliferation in human tissues is the unavailability of two key pieces of information: the quantification of DNA amount and measurement of active DNA replication. DNA staining by intercalating agents, such as Hoechst, is routinely performed in tissue imaging, and the cyclic immunofluorescence methods rely on it to align the rounds of imaging to obtain highly multiplexed single-cell datasets. However, the DNA quantification from tissue images does not appear to be an accurate representation of DNA content. We found that single cell distributions of DNA content from tissue are not bimodal (Figure S7), unlike those in cell culture (Figure S3I). In addition, genomic instability and aneuploidy are a common feature in human tumors, making the interpretation of DNA quantification in solid tumors additionally challenging. Similarly, the detection of DNA replication - usually performed by EdU incorporation in cell culture - is not technically possible in fixed human tissues.

**Supplemental Note S4: Limitations of multichannel gating to infer cell cycle distributions**

In order to attempt to understand the cell cycle position of cells based on a gating approach, we gated single channels and looked at the combinatorial patterns of positivity for multiple channels, or “multichannel gating” (Figure S3D). Binarizing an 8 cell cycle marker panel by gating produces 2^8^ = 256 combinatorial states. In Figure S3D we show a visual representation of the top 50 most frequently occurring states using same breast tumor tissue that was previously used for the single distribution and two-channel scatters in Figure S3D). We used the Upset plot package (https://gehlenborglab.shinyapps.io/upsetr/) to display the combination of positive markers in each of the 50 states and the relative abundance of each state. While the abundance of these states is technically quantifiable, it is important to remember that in the absence of bimodal single marker distributions, the choice of gating threshold is somewhat arbitrary. Small changes in any of the cutoffs would drastically modify the relative abundances of all the states.

In the absence of information about the ploidy and DNA replication state of the cells, discriminating the combinatorial states generated by multichannel gating is prohibitive. In cell culture experiments, both DNA amount and EdU incorporation have bimodal distribution, the combination of which clearly defines the G1 (2N, EdU-), S (EdU+) and G2/M (4N, EdU-) phases of the cell cycle. This information is instrumental to define cell cycle states, because the dynamics of cyclin expression (as well as CDT1, Geminin and phospho-Rb) are much more graded through the cell cycle. In addition, while the dynamics of single proteins through the cell cycle are well-characterized, it is still unclear how they relate to each other. For example, the levels of Cyclin D1 decrease from G1 to S/G2, but how does this quantitatively relate to the increase in Geminin in tissue? Similarly, how does the rise of G2-phase cyclins like Cyclin A1/2 relate to the dynamics of decrease in CDT1? Without firm answers to these questions we are not able classify the multichannel gating data in Figure S3D.

Overall, the loss of information caused by the gating binarization of the continuous distributions did not simplify the classification problem; it only changed the nature of the problem from continuous to discrete. Thus, because of 1) the lack of interpretability and 2) the lack of robustness in state abundances, we concluded that a gating strategy would not lead to insights in the cell cycle analysis (specifically in our multiplexed imaging datasets). Instead, we pivoted to methods that use the full continuum of the markers’ expression values to create unsupervised metrics for cell cycle analysis.

**Supplemental Note S5: Robustness of the ccD-CMD algorithm and coherence metrics**

The ccD-CMD algorithm does not have a stochastic component and hence it is an injective function with one-on-one mapping from input to output. However, its application to estimate the coherence metrics IOV and CFD is sensitive to two variables: 1) the number of cells from the tissue used in input, 2) the markers used to calculate the cell cycle difference (ccD).

Because the ccD is a quantification of the binary distances between pairs of cells, the computing time to calculate the ccD increases non-linearly with the number of cells provided in input. It hence becomes prohibitive to run the ccD-CMD computation on more than a few tens of thousands of cells. On the other hand, when using tissue microarray cores, the number of cells available for analysis is limited. For both these reasons it is important to estimate how sensitive the performance of the algorithm is to the number of cells in input. In Figure S4D we calculated the CFD and IOV parameters in 5 tissues using an increasing number of cells (“n” from 50 to 2000, breast cancer tissue ROIs from Figure 4F with at least 20,000 MPI+1 cells were used for this analysis). For each n, we run the ccD-CMD algorithm 40 times with a different set of n cells from the same tissue and calculate the coefficient of variation (CV) for the IOV and CFD metrics. For both metrics the CV quickly decreased and reached a plateau. This shows that for whole tissue experiments using more than 2000 cells would not provide a great increase in precision. In addition, when cell numbers are limited, using at least 500-1000 cells is required for an acceptable estimate.

Further, we tested the dependency on single markers on the coherence metrics, by eliminating one or two markers from ccD calculation (Figure S4E). For this we used the MCF10A cell line dataset, for which we have experimental controls obtained by perturbing the cell cycle with palbociclib and nocodazole (Figures 3G-3J). Eliminating one marker (out of 10 in the panel) had limited effect, comparable to the biologic variability of the three replicates in Figure 3J. Eliminating two markers did not produce much larger displacement from the whole-panel estimates (green dot) than the single marker removal. Furthermore, the direction of the changes is spread in all directions of the IOV-CFD phase plane, suggesting that addition of markers converges to a central estimate, the green dot. In conclusion, these results show that the ccD-CMD algorithm is robust to the removal of one or two markers, especially when compared to experimental perturbations.

**Supplemental Note S6: Comparison between ccD-CMD and published time inference methods.**

Many algorithms have been published to computationally infer ordering of cells based multidimensional data but most of these algorithms have been developed to analyze genomics datasets, rather than proteomics or imaging ones. We selected a subset of these time inference algorithms, processed our multiplexing imaging data and compared the results with the ccD-CMD representation and pseudotime ordering from the same datasets. We used the following three algorithms, all originally developed to process single cell RNA sequencing data: SCORPIUS (Cannoodt et al., 2016), Palantir (Setty et al., 2019) and Cyclum (Liang et al., 2020).

We compared the ccD-CMD time inference output with SCORPIUS, Palantir and Cyclum on three exemplar CyCIF datasets from different experimental sources:

1. on tissue-based CyCIF data from a breast tumor tissue sample from Figures 3A-3F;
2. on plate-based CyCIF data from unperturbed MCF10a cells;
3. on synthetic data generated from a mathematical model of the mammalian cell cycle.

The results of the comparisons are shown in Figures S3H and S4A-S4C. Overall, we found that the results from Palantir and Cyclum did not recapitulate the cell cycle dynamics in any of the experimental settings. Palantir is designed to model cell differentiation into terminal states, which does not describe the trajectory of cell cycle dynamics. Cyclum uses an autoencoder with non-linear periodic transformation functions to infer a latent circular trajectory. Although Cyclum is designed for cell cycle dynamics, it does not appear to be able to detect any discernible structure in multiplexed imaging data. Cyclum’s expected input is scRNA-seq data, which has a) three orders of magnitude more features than our datasets, and b) extremely different type of noise than imaging data. Additionally, because Palantir and Cyclum use non-linear dimensionality reduction methods, they are significantly slower than CMD scaling, which makes them less suitable for the high-throughput analysis of large datasets.

The time ordering output from SCORPIUS closely resembles the ccD-CMD output. However, the ccD-CMD reduced dimensionality representation is strikingly different from the SCORPIUS output. While the data points in the ccD-CMD representation formed a torus, which allowed us to parametrize the representation and derive the “cell cycle coherence” metrics above, SCORPIUS’s cloud output could not be similarly parametrized. In the tissue data (Figure S4A), the ordering by Cyclum and Palantir does not capture the basic tenets of cell cycle protein dynamics. In addition, the t-SNE representation of the Palantir output shows a branched structure, which is incompatible with the cell cycle. These observations are true for all the datasets we describe below. SCORPIUS and the ccD-CMD ordering is comparable, but a directly comparison is not possible as the ground truth cell cycle ordering in tissues is unknown. In the cultured cell line data (Figure S4B), the ccD-CMD cell ordering appears comparable to the SCORPIUS ordering when run with single cell data from unperturbed cells (MCF10A). In both orderings, there is a clear inverse relationship between CDT1 and Geminin as well increasing DNA content as the cell cycle progresses. However, many of the markers show the drawbacks of SCORPIUS’s non-cyclical path. For example, Geminin, phosphor-RB, and Cyclin B show a rise from low to high through time but do not show the drop from high to low that is expected in the cyclical pattern of cell cycle dynamics. ccD-CMD is able to better capture this cyclical pattern. To obtain a dataset with a ground truth for cell ordering, we simulated the cell cycle protein dynamics with a system of ordinary differential equations (ODEs) based on the model by Csikasz-Nagy and Tyson (Figure S3E, Csikász-Nagy et al., 2006) and generated values of nine cell cycle markers over time. We ran both ccD-CMD and SCORPIUS and compared the reconstructed orderings with the known ordering from the differential equation numerical solution. ccD-CMD outperformed SCORPIUS. In the ccD-CMD ordering, 93% of cells were within 1% of their correct ordering as opposed to 36% of cells in the SCORPIUS ordering. It is notable that the two-dimensional representation generated from ccD-CMD has cells tightly distributed along the circle with the exception of a gap where M phase is expected to be (Figure S3F). The ccD-CMD ordering also shows the most misordering in M phase/early G1 (Figures S3G and S3H), which reflects the difficulty of detecting and ordering cells in M phase that is also seen in the multiplexed imaging data.

These comparisons show that the ccD-CMD algorithm orders cells efficiently and that the resulting dynamics are congruent with literature on cell cycle biology. Both Palantir and Cyclum do not seem able to recapitulate the basic tenets of cell cycle protein dynamics, especially in the tissue exemplar dataset. Time inference obtained using SCORPIUS and the ccD-CMD is comparable. In experimental data (either cell line or tissue data), the ground truth information is not available, and hence quantifying the improved precision of ccD-CMD over SCORPIUS is not possible. However, on synthetic data ccD-CMD has a higher ordering precision compared to SCORPIUS. Finally, the main advantage of the ccD-CMD representation over the other time inference algorithms is the ability to provide a reduced-dimensional representation of “coherence” in cell cycle dynamics. The ccD-CMD representation in control systems forms the torus-shaped that inspired the circle fit approximation. Other algorithms’ reduced representation did not show an actionable topology (Figures S4A and S4B), either because of a lack of reduced dimension representation (Cyclum), a lack of topological shape (SCORPIUS), or a disconnected and branched topology (Palantir). In tissues we detected quantifiable differences in the ccD-CMD representation topology across patients, and we chose three patient samples to portray the range of topologies observed (Figure 4C). However, the SCORPIUS and Palantir outputs from the same data show no discernible difference between the three datasets (Figure S4C).

## SUPPLEMENTAL EXPERIMENTAL PROCEDURES

### Pairwise cell cycle difference and classical multidimensional scaling (ccD-CMD)

#### Description of input

For this paper, we constructed trajectories of cell cycle progression from single cell immunofluorescence measurements obtained from fixed tissue images. We utilized the MPI classification to isolate epithelial proliferating cells (i.e., MPI +1 cells) and used only those as input. For the cell culture experiments all cells measured were used in input, because the cell composition is homogeneous and the majority of cells were MPI +1. Each cell was represented by a cell cycle marker vector of normalized values for markers of cell cycle proliferation (Ki-67, PCNA, MCM2), cell cycle arrest (p21, p27), and cell cycle progression (phospho-RB Ser807/811, CDT1, Geminin, and Cyclins A1/2, B1, D1, and E1).

The markers for the ccD-CMD analysis were chosen to represent multiple cell cycle transitions. For instance, p21, CDT1 and Cyclin D1 cover the transition through G1 and into S-phase. phospho-RB, Ki-67, Cyclin A1/2 and B1 cover the passage from S-phase into early and then late G2 phase. Although PCNA and MCM2 are used to calculate the MPI, they were not largely used in the ccD-CMD trajectory inference because their variability within MPI+1 cells is minimal; therefore, they do not provide additional information and potentially add spurious noise. This reasoning also holds true for markers that are highly specific, but are only expressed in a small subset of cells (e.g., phosphor histone H3). It’s important to note that not all of these markers were usable for every dataset, especially in tissue-based experiments where the staining variability can be high and where individual tumors might genetically or epigenetically lose the expression of single proteins (see Table S2 for details of which markers were used for the ccD-CMD calculations in each experiment).

#### Dimensionality reduction

The ccD-CMD starts by calculating the cell cycle difference (ccD) matrix, which is defined as the absolute value of the pairwise Pearson correlation between the cell cycle marker vector of normalized values of each cell. An example of a ccD matrix is presented in Figure 3C.

In order to be interpreted the ccD matrix needs to be reduced in dimensions. Classical multidimensional scaling (CMD) is a linear dimensionality reduction method used to approximate the pairwise distances between n points (in our case n = number of single cells) to a representation in lower dimensions. Commonly, reduction to two dimensions is chosen to ease visualization and interpretation. CMD scaling is a linear algorithm, and although assuming linearity is an oversimplification, CMD scaling runs significantly faster than non-linear methods, which is important for scalability as we routinely run the algorithm on hundreds of samples with upwards of 20,000 cells.

#### Circular fitting: trajectory model and cell ordering

The reduced two-dimension ccD scatter, referred to as the “ccD-CMD” representation, or landscape, is parametrized by fitting it to a circle by least-squares minimization. This choice was made with the observation that the two-dimensional representation has an underlying cyclical structure following the dynamics of the cell cycle. This is in contrast with many other time inference algorithms that search for non-cyclical paths through the data.

The circular fitting served two purposes:

1. to perform ordering of the cells around the cell cycle position. Notably the ordering is only most accurate for populations of cells where the ccD-CMD representation forms an evenly distributed torus (referred to as “cell cycle coherent” in the main text), rather than a skewed torus or amorphous point cloud (referred to respectively as “skewed cell cycle” and “non-canonical”).
2. to parametrize the ccD-CMD representation and extract quantitative metrics that summarize the sample’s overall cell cycle temporal organization.

For each data point (i.e., single cell), two parameters are calculated, 1) the shortest distance between the data point and the fitted circle (*d*) and 2) the angle relative to the point of origin of the fitted circle (θ) as shown in Figure 3E. Using the angle, each data point was projected to the nearest point on the c^θ^ircle to order the cells in what is referred to as the “cell cycle ordering.” Given the cyclical nature of the ordering, the point of origin (time = 0 or start time) is arbitrary. For each population of cells being analyzed, two overall metrics were extracted from the fit:

1. the circle fit distance (CFD), which is the mean *d* of the population
2. the inter-octile variation in angle (IOV). To calculate the IOV, the angle measurements were binned to 8 equally sized bins and the proportion of cells in each bin were calculated. To ensure lack of positional bias, the proportion calculation was repeated by shifting the bin position by π/8. The IOV is the coefficient of variation of the bin proportions, which is equal to 0 in a uniform^π^ly distributed population.

#### Measures of cell cycle coherence

The two metrics, CFD and IOV, were used to describe what we call the “cell cycle coherence” of a sampled cell population. A low CFD indicates that the points in CMD space are distributed along the circular path (best-fit circle). As more and more cells accumulate in the center of the ccD-CMD representation (forming a cloud of points) the CFD metric increases. In the ccD-CMD representation, the 2D distance between two points is a lower-dimensional approximation of their difference in multidimensional space, in our case the cell cycle space. Therefore, a cloud of points means that the cell cycle marker vectors of cells in the cloud are similar, because the difference between cell cycle states is not sufficiently distinct to separate them in ccD-CMD projection i.e. there is no coordination between cell cycle markers (e.g., Sample 3, Figure 4C). If the measured cells are random samples from a deterministically oscillating system, the distance between the cells would be proportional to the time between the positions of the random samplings. In this case, the data points would form a topological toroidal shape and in turn, a low CFD. In Figure 5, we showed how a sharp drop in HER2 expression led to a shift from a toroidal ccD-CMD representation (9 weeks HER2 “on”) to a point cloud (2 days HER2 “off”), with a corresponding increase in CFD (from 30 to 50). Our interpretation is that HER2 expression promotes a strong signal for cells to grow; once this is withdrawn, the system drifts in multiple directions and the levels of the cell cycle markers lose coordination.

A low IOV indicates that points are distributed evenly around the circle as opposed to clustered at specific regions. This clustering occurs if cells slow down in a specific part of the cell cycle, hence accumulating at that location. Notably, a similar clustering would occur if cells were to accelerate through a specific part of the cell cycle, but in this case, the accumulation of cells would occur away from the acceleration point. In both instances, the population of cells would be unevenly distributed and have high IOV, a state that we call “skewed cell cycle” because of the uneven distribution of cells around the ccD-CMD torus. The extreme case scenario is cell cycle arrest, for instance triggered in MCF10A by cell cycle inhibitors, palbociclib and nocodazole (Figures 3G-3J). In both treatments, at both 24 and 48 hours the recorded IOV is up to three-fold higher than freely cycling MCF10A cells. However, these perturbations are extreme scenarios. In human samples, we focus on the MPI +1 population of cells and eliminate from the analysis the most likely arrested population (MPI -1, with high levels of p21 or p27). Accordingly, most human tumors we measured had IOV between 0.4 and 1.2, while the fully arrested cells have IOV of 1.5-2. The high IOV state potentially represents populations of cells moving towards cell cycle arrest. However, as described above, high IOV could also suggest a more streamlined cell cycle with a shortening in one cell cycle phase and in fact a faster, albeit still less balanced, completion of the cell cycle.

---

**Figure S1.**
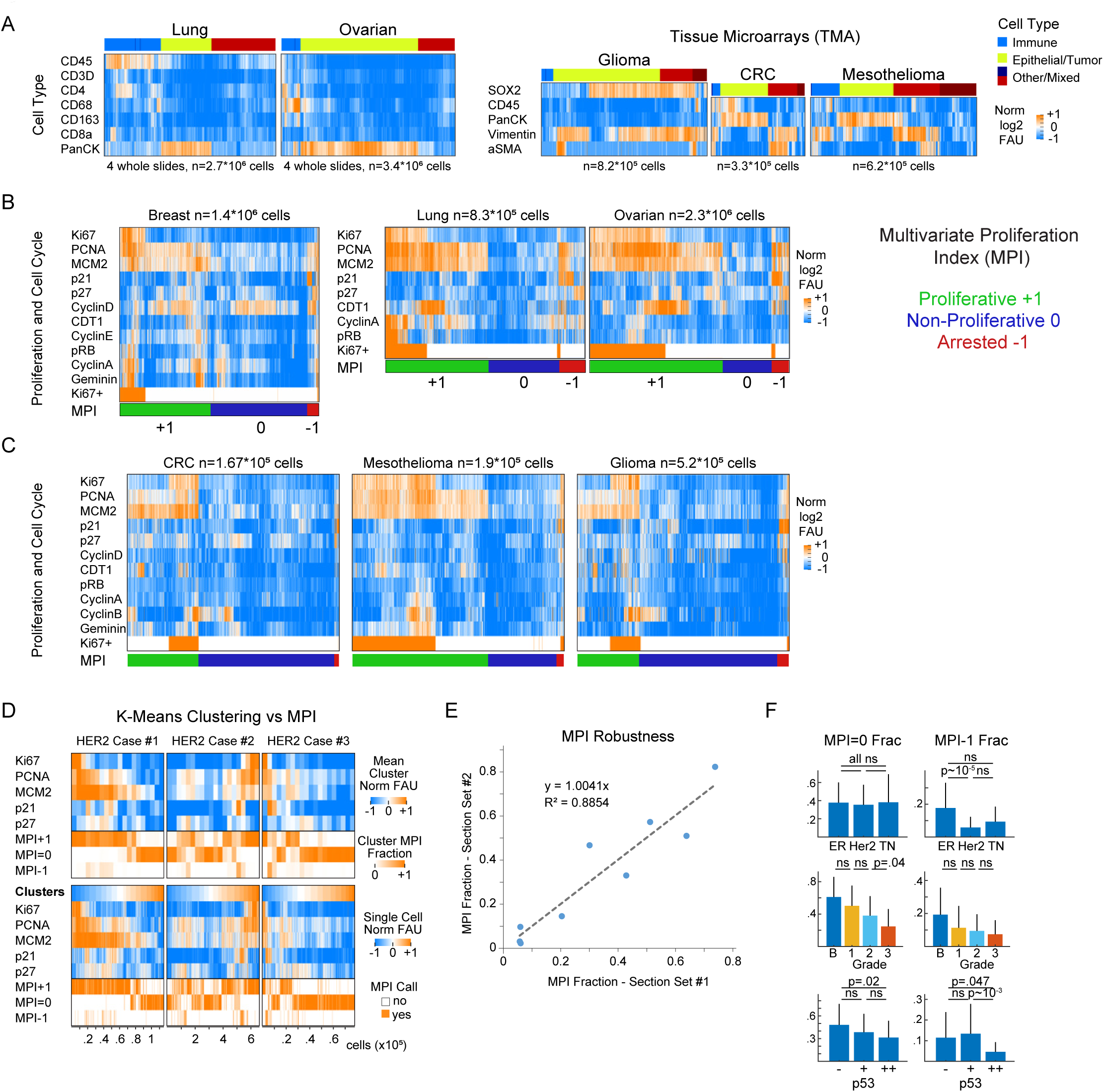
Multivariate Proliferation Index (MPI) in human cancer tissues. (A) Clustered heat map of log2 normalized cell lineage marker signal intensities on a per-cell basis derived from t-CyCIF images of 4 whole slides of lung squamous cell carcinoma (SCC) and ovarian carcinoma, and tissue microarrays (TMAs) from glioma, colorectal carcinoma (CRC) and mesothelioma. (B-C) Clustered heat map of single-cell signal intensities of cell cycle markers for epithelial/tumor cells in Figure 1A (breast carcinoma) and panel S1A above. Ki-67+ cells were identified by normalization using Gaussian mixture modeling with 2 components. Multivariate Proliferative Index (MPI) indicated: +1 (proliferative, green), 0 (non-proliferative, blue), or -1 (arrested, red). (D) K-means clustering heat map of five MPI markers for epithelial/tumor cells of three HER2+ breast cancer samples (k = 20 clusters), and heat map of single-cell normalized log2 intensities. In both, the corresponding MPI category is depicted for comparison (in k-mean clustering fraction of MPI category is depicted). (E) MPI robustness comparison between two sets of serially cut tissue sections on the 3 breast HER2-positive cases from Figure 1. Each dot is the fraction of cells in one MPI category in both sets of tissue section (linear least-square fit with fixed origin at y=x=0). (F) Comparison of MPI 0, and MPI -1 fractions in epithelial/tumor cells across different classifiers of breast cancer in Figure 1D (KS p-values with 0.05 significance cutoff).

**Figure S2.**
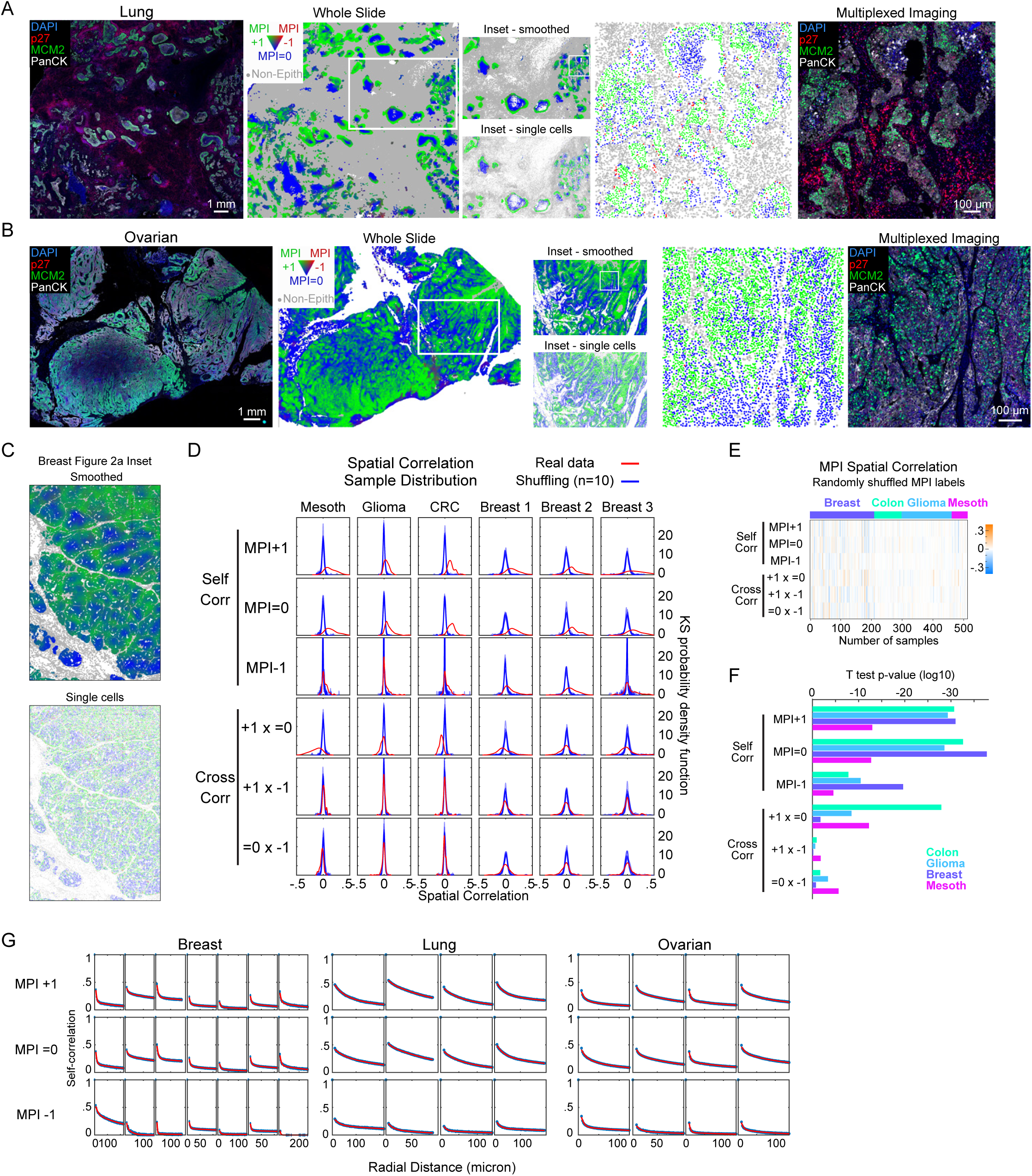
Multivariate proliferation index (MPI) short- and long-range correlations in human cancer tissues. (A-B) Composite t-CyCIF image from (A) lung squamous cell carcinoma, and (B) ovarian carcinoma (scale bar 1mm) and corresponding image of long-range spatial maps of MPI categories (smoothed over 40 neighboring cells for visualization purposes only). Inset panel showing both smoothed and single-cell MPI calling. Further inset panel of single-cell MPI calling and corresponding composite t-CyCIF image (scale bar 100 µM, white = pan-cytokeratin, green = MCM2, red = p27, blue = DNA). (C) Spatial maps of MPI categories smoothed over 40 neighboring cells and non-smoothed (single-cell calling) for inset from Figure 2A. (D) Red lines, sample distribution of spatial correlations within and across (“self corr” and “cross corr”) MPI categories for mesothelioma, glioma, colorectal carcinoma (CRC) and three breast tissue microarrays (n = 52, 163, 89, 69, 85, 57 samples respectively, k = 5^th^ neighbor approximation, KS density approximation). Blue lines, bootstrap distribution comparison obtained by randomly shuffling of MPI labels (10 independent shuffles). (E) Heat map of spatial correlations within and across randomly shuffled MPI labels (“self corr” and “cross corr”, k = 5^th^ neighbor, n = 513 samples). (F) t-test p-values for red line distributions in panel D (log10 scale used for visualization purposes). (G) Spatial correlation plots with two exponential fit for the three MPI categories across breast, lung, and ovarian tumors (whole slide imaging, Figure 2D). Each column represents an individual and independent tissue.

**Figure S3.**
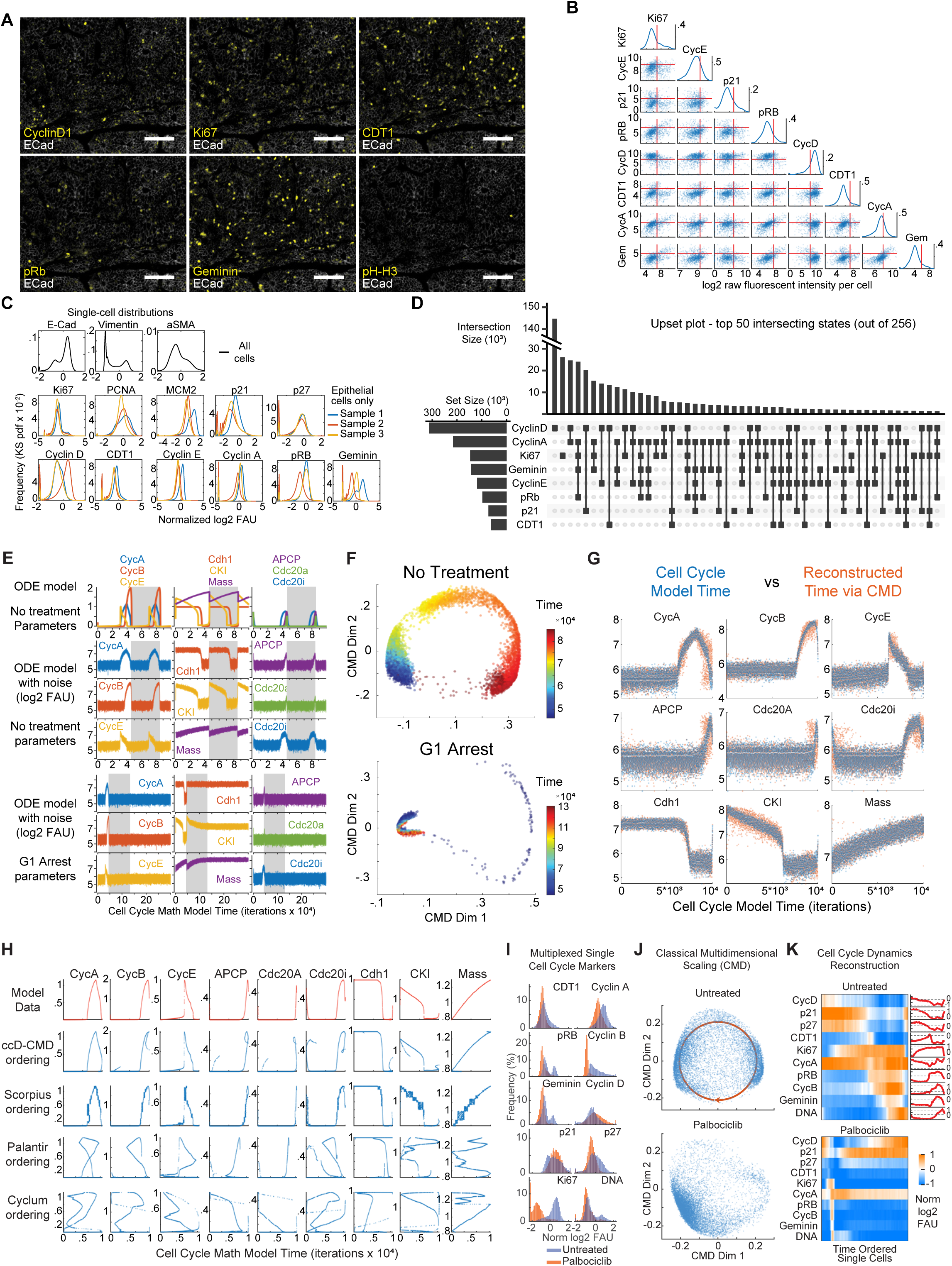
Cell cycle marker single cell distributions, multi-channel gating and testing of ccD-CMD framework. (A) Representative single-channel cell cycle marker images from t-CyCIF imaging with e-cadherin from tissue in Figure 3A (scalebar 100 µm). (B) Single-channel distributions and two-dimensional scatter plots of a subset of cell cycle markers from HER2 positive breast cancer tissue case 2 in Figures 1C and 1E (n = 1,000 epithelial/tumor cells, log2 signal per cell prior to normalization). (C) Single channel distributions for cell lineage, proliferation, and cell cycle markers from three HER2+ breast cancer samples (KS density approximation). For proliferation and cell cycle markers only epithelial/tumor cells were used. (D) Upset plot of three HER2+ breast cancer samples (n = 0.52 million cells) showing frequency of cell cycle marker positivity and their combinations sorted by frequency. (E) Traces from generalized model of mammalian cell cycle (Csikász-Nagy et al., 2006). Top panels, no noise added. Middle panels, Gaussian measurement noise added (additive and multiplicative). Bottom panels, simulation switch to G1 arrest parameters (CDK/CyclinD complex set to 0) after one cell cycle. Grey areas are the time frame used for plots in panels F and G. (F) Classical multidimensional scaling of cell cycle difference (ccD-CMD) of mathematical model results in shaded areas of panel E. Color is the time variable in the mathematical model (n = 10,000 points). (G) Comparison of model’s cell cycle time (blue dots) and time reconstructed (orange dots) by fitting a circle to the ccD-CMD plot in panel F. (H) Comparison of time inference methods ccD-CMD, SCORPIUS, Palantir and Cyclum applied to synthetic data generated by the mathematical model in panel E (n = 10,000 observations). (I) Histograms of the fluorescence signal of cell cycle markers measured at single-cell level using plate-based CyCIF from untreated (blue) and 24 hours palbociclib treated (orange) MCF10A cells grown in culture (n = 10,000 cells per condition). (J) Classical multidimensional scaling of ccD reduced to 2 dimensions. Scatter plot for cells in panel I (red, circle fit, n = 10,000 cells per condition). (K) Heat map and time plot of single-cell signal intensity measurements of cell cycle markers ordered from left to right in cell cycle time (normalized log2 fluorescent arbitrary units moving mean over 200 cells).

**Figure S4.**
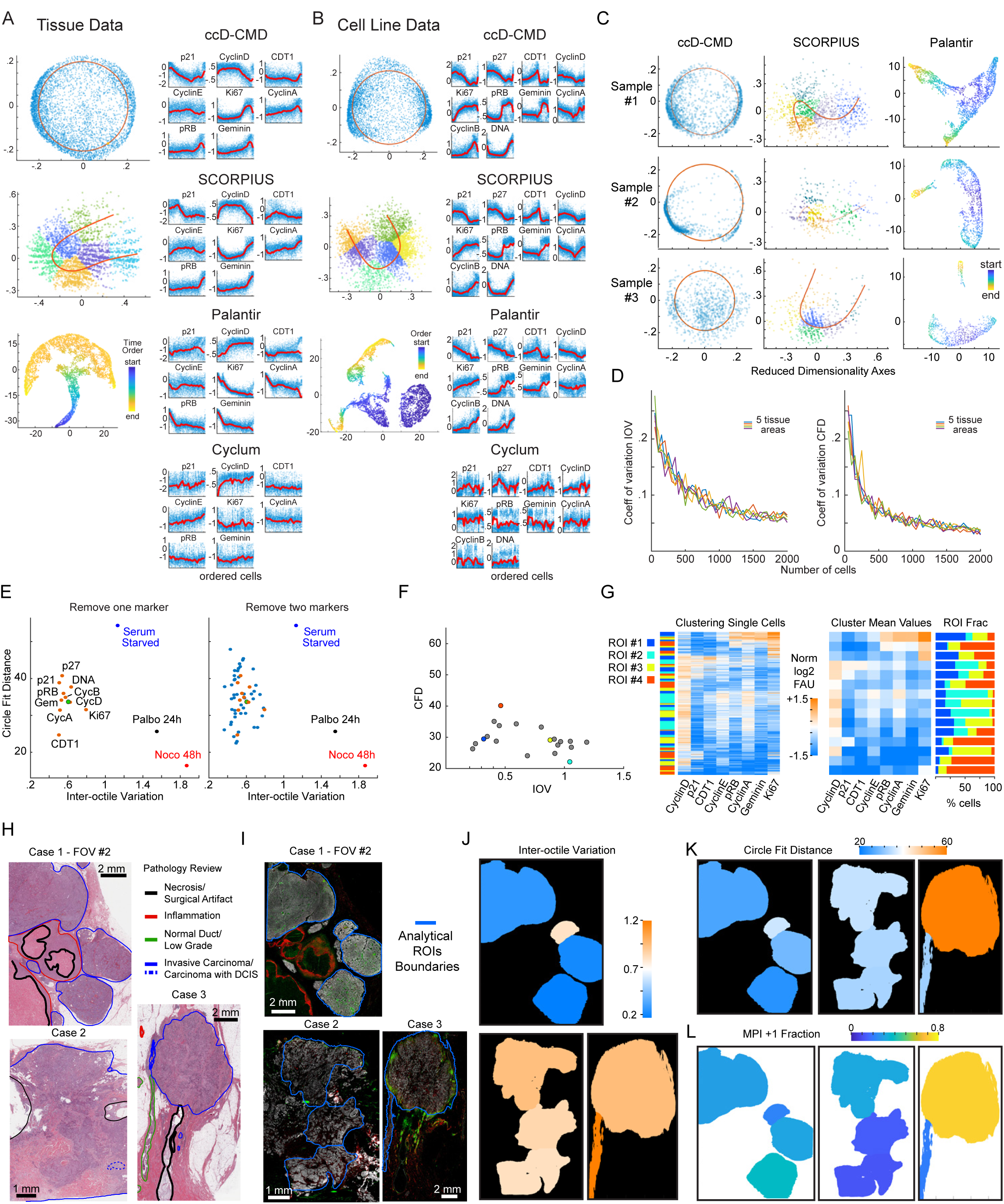
Temporal inference of cell cycle dynamics from ccD-CMD from human cancer tissues. (A-B) Comparison of time inference methods ccD-CMD, SCORPIUS, Palantir and Cyclum from (A) HER2+ breast cancer tissue data from Figures 3A**-3F** and (B) MCF10A untreated cells from Figure 3G (n = 5,000 cells, same cells used for all algorithms). Left, two dimensional visualization output. Right, pseudotime ordering output (normalized log2 fluorescent arbitrary units, moving mean with 200 cells window). (C) Comparison between two dimensional reduced space visualization from three time inference algorithms with data from HER2+ breast patient samples 1,2, and 3 from Figures 4B**-4D**. (D) Coefficient of variation of Inter-Octile angular Variation (IOV) and circle fit distance (CFD) in 5 tissues using increasing number of cells (n > 20,000 cells per tissue, CV calculated over 40 independent sub-samplings). (E) Comparison of coherence metrics IOV and CFD when one or two markers are removed from ccD-CMD algorithm. Data from untreated MCF10A cells used in panel b. Green dot is original representation. Orange, one marker removed. Blue, two markers removed. (F) Scatterplot of Circle Fit Distance (CFD) against Inter-Octile angular variation (IOV) for each ROI from Figure 4F. (G) K-means clustering of cell cycle markers from selected ROIs used in Figure 4G (n = 3600 cells per ROI, k = 15 clusters). Left, single cell clustering with ROI annotation (log2 normalized FAU). Middle, cluster median. Right, ROI composition for each cluster. (H) Scanned image of hematoxylin and eosin (H&E) stained section from three HER2 positive breast tissues with pathology annotations. (I) Composite t-CyCIF images of fluorescence of tissues from panel H. (J) Inter-Octile Variation IOV, (K) Circle Fit Distance (CFD), and (L) MPI +1 fraction for each ROI noted in panel I.

**Figure S5.**
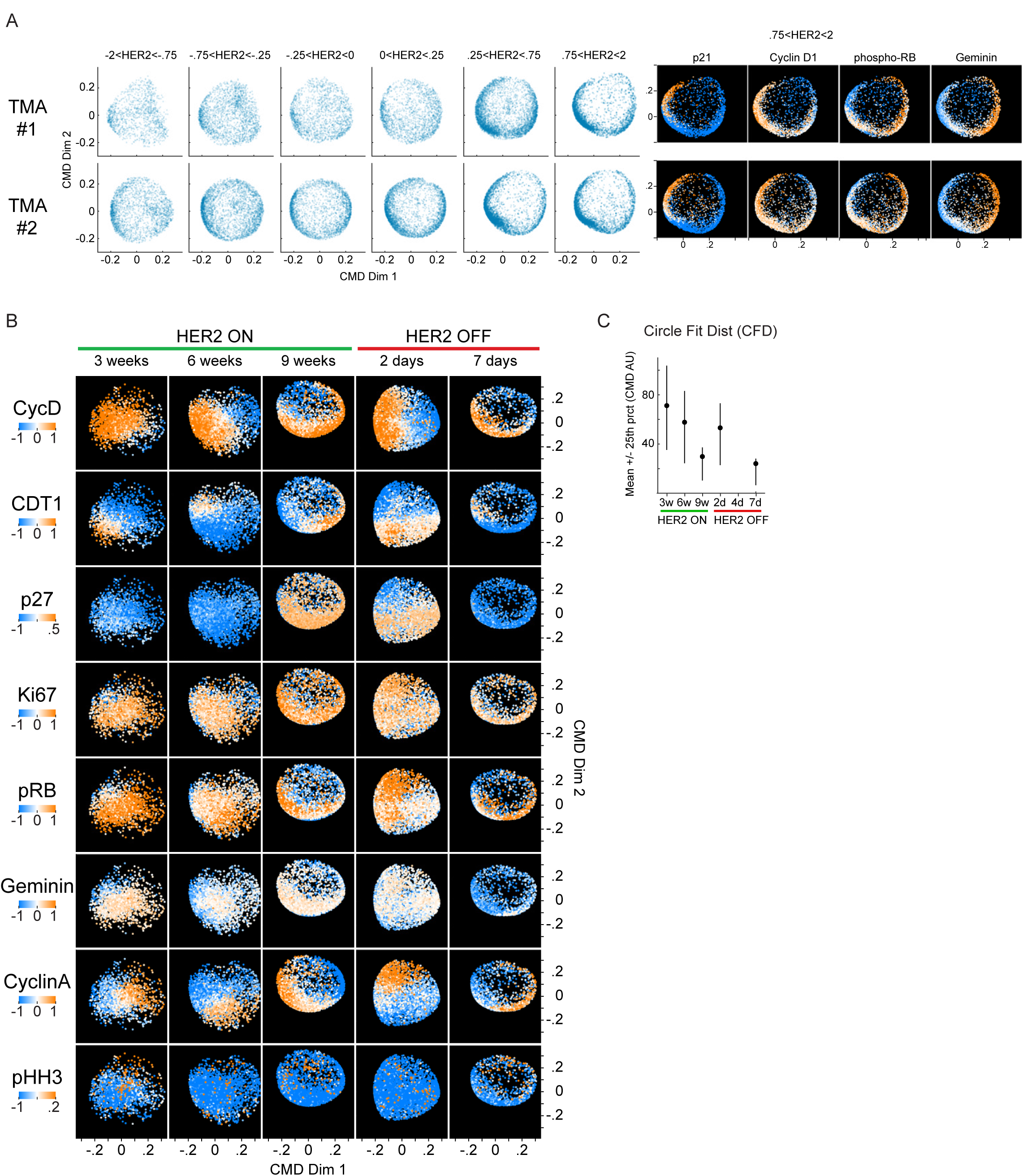
HER2 expression and cell cycle dynamics in human breast cancer tissues and HER2 driven mouse model of breast cancer. (A) ccD-CMD plots for two HER2+ breast cancer tissue microarrays (TMA # 1 and #2) for increasing levels for HER2 protein. Single cells were binned by normalized HER2 levels (n = 5,000 cells per bin were used). Right, ccD-CMD scatter plots of highest HER2 bin for both TMA1 and 2 with single marker normalized intensities mapped to color (n = 5,000 cells). Left, single marker normalized intensities mapped to color for highest HER2 bin. (B) ccD-CMD scatter plots of the single-cell data from MPI +1 cells from time course of HER2 induction and repression in GEMM with single marker normalized intensities mapped to color (n = 5,000 cells per plot, p27 was not used by the ccD-CMD algorithm). (C) Mean +/- 25^th^ percentile of circle fit distance of cell cycle markers in tumor cells (in situ and invasive) from Figure 5H.

**Figure S6.**
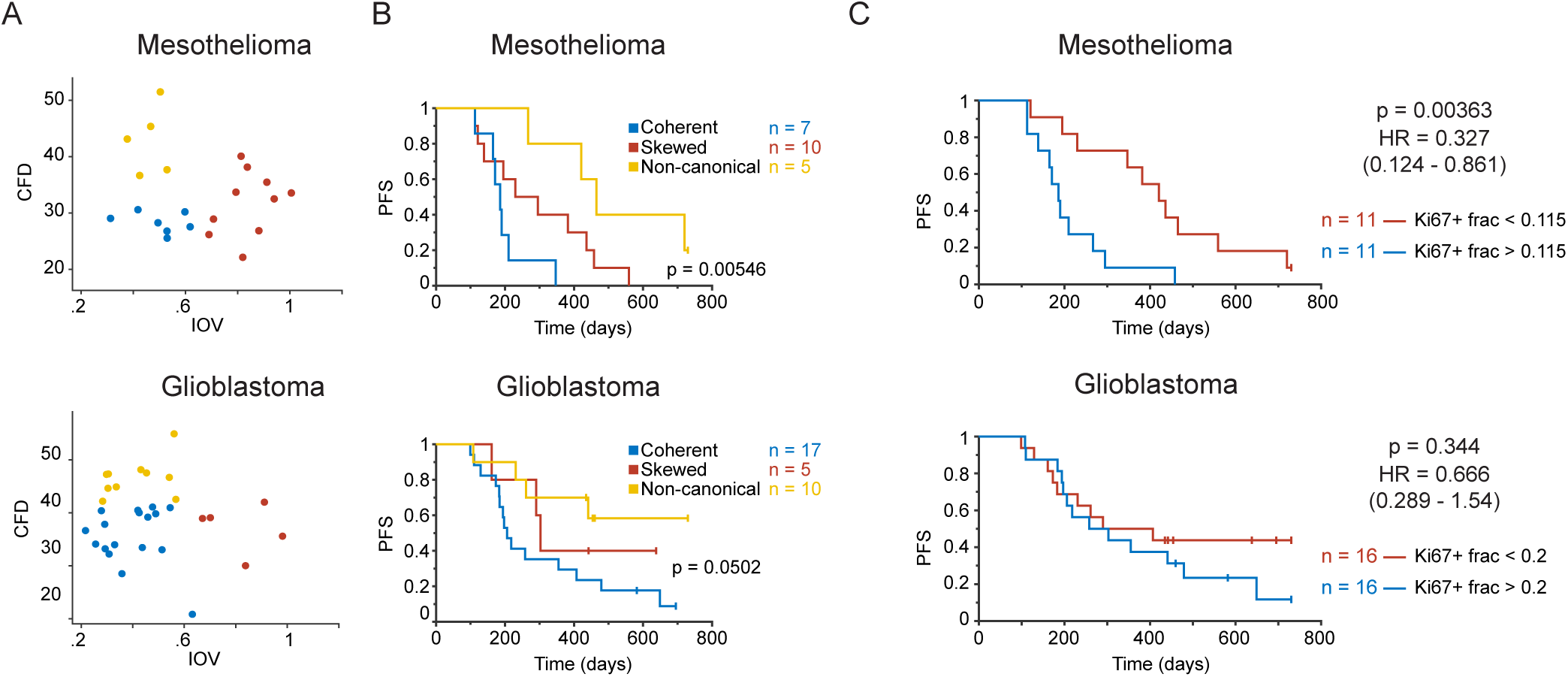
Association between with clinical outcome and cell proliferation metrics. (A) Scatterplot of CFD versus IOV from a mesothelioma and glioblastoma sample cohorts (n = 22 and 32 patients). Colors represent binning into coherence groups according IOV and CFD metrics. (B-C) Kaplan Meier estimation and logrank p-value of progression-free survival (PFS) for the two patient cohorts in Figure 7. (B) Patients binned in three groups “coherent”, IOV^high^ “skewed” and IOV^low^ CFD^high^ “non-canonical” groups from panel A). (C) Patients binned in two groups based on the median Ki-67+ fraction.

**Figure S7.**
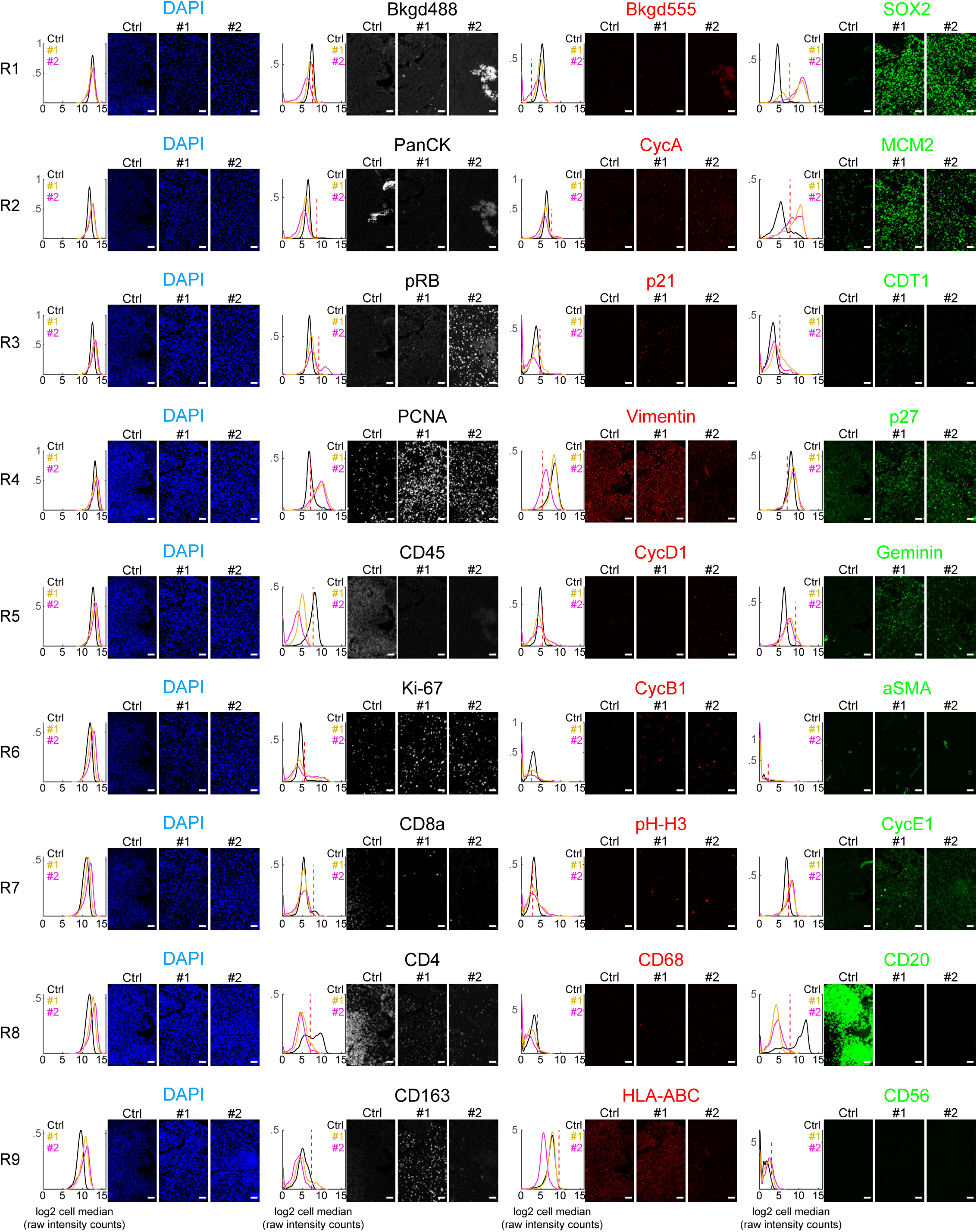
Example of 27-plex t-CyCIF experiment with tumor sample and tonsil control tissues. Example of positive and negative staining for all markers in t-CyCIF experiment through rounds of cyclic imaging. Three independent samples are shown, Ctrl is a non-malignant tonsil tissue sample, #1 and #2 are glioma samples. Plots are single-cell kernel density estimation for patient samples from respective images (median per pixels within the cell area, log2 FAU, not normalized, black=control tonsil, yellow = sample #1, magenta = sample #2, n = 4278, 2629 and 2609 cells respectively). Each row of images and data is a successive round of t-CyCIF in the same tissue area (Rx is the x^th^ round of imaging). All images from antibody channels were linearly contrasted between 0 and 2000 fluorescence units for ease of comparison. Scale bar 50 µm.

